# A lipopolysaccharide-dependent phage infects a pseudomonad phytopathogen and can evolve to evade phage resistance

**DOI:** 10.1101/2022.06.10.495718

**Authors:** Suzanne L. Warring, Lucia M. Malone, Jay Jayaraman, Richard A. Easingwood, Luciano A. Rigano, Rebekah A. Frampton, Sandra B. Visnovsky, Shea M. Addison, Loreto Hernandez, Andrew R. Pitman, Elena Lopez Acedo, Torsten Kleffmann, Matthew D. Templeton, Mihnea Bostina, Peter C. Fineran

## Abstract

Bacterial pathogens are major causes of crop diseases, leading to significant production losses. For instance, kiwifruit canker, caused by the phytopathogen *Pseudomonas syringae* pv. *actinidiae* (*Psa*), has posed a global challenge to kiwifruit production. Treatment with copper and antibiotics, whilst initially effective, is leading to the rise of bacterial resistance, requiring new biocontrol approaches. Previously, we isolated a group of closely related *Psa* phages with biocontrol potential, which represent environmentally sustainable antimicrobials. However, their deployment as antimicrobials requires further insight into their properties and infection strategy. Here, we provide an in-depth examination of the genome of ΦPsa374-like phages and show that they use lipopolysaccharides (LPS) as their main receptor. Through proteomics and cryo-electron microscopy of ΦPsa374, we revealed the structural proteome and that this phage possess a T=9 capsid triangulation, unusual for myoviruses. Furthermore, we show that ΦPsa374 phage resistance arises *in planta* through mutations in a glycosyltransferase involved in LPS synthesis. Lastly, through *in vitro* evolution experiments we showed that phage-resistance is overcome by mutations in a tail fiber and structural protein of unknown function in ΦPsa374. This study provides new insight into the properties of ΦPsa374-like phages that informs their use as antimicrobials against *Psa*.

**Originality-Significance Statement:** The rise of phytopathogen resistance to agrichemicals poses a significant threat to crop production, and requires urgent attention. The work presented here examines a phage genus, members of which utilize LPS as a receptor, and show potential as biocontrol agents of *Psa*. Studies *in planta* showed the development of *Psa* resistance to the phage, whilst maintaining bacterial virulence. This finding underscores the importance of examining phages in their ecological context for the informed design of phage cocktails that are effective in phytopathogen control.

## Introduction

*Pseudomonas syringae* pv. *actinidiae* (*Psa*) is the causal agent of bacterial canker in kiwifruit, and has caused a significant economic disruption to the global kiwifruit industry (Takikawa *et al*., 1989; Balestra *et al*., 2009, 2010; Ferrante and Scortichini, 2009; Koh *et al*., 2010; Vanneste *et al*., 2011; Bastas and Karakaya, 2012). Treatment of infected orchards and control of the disease has been achieved by copper-based agrichemicals, antibiotics, removal of infected plants and replacement of susceptible kiwifruit cultivars with more resilient ones. However, recent whole genome sequencing of orchard isolates has revealed the emergence of both copper and antibiotic resistance (Cameron and Sarojini, 2014; Colombi *et al*., 2017). This threat, coupled with environmental concerns surrounding the use of copper and antibiotics in food production, has created the need for alternative methods to control *Psa*, and bacteriophages may offer a solution (Buttimer *et al*., 2017; Dy *et al*., 2018).

Previously we isolated a library of phages capable of infecting pandemic *Psa* variants, which therefore represented potential biocontrol agents (Frampton *et al*., 2014). Further studies from our group and others have characterized various phages with biocontrol potential against *Psa* (Frampton *et al*., 2015; Yu *et al*., 2015; Park *et al*., 2018; Wojtus *et al*., 2019; Yin *et al*., 2019; Martino *et al*., 2021). While these studies focused on basic phage characterization, recent work has illustrated the potential efficacy of phage treatment of *Psa*-infected kiwifruit plants (Flores *et al*., 2020; Song *et al*., 2021). Although such research illustrates the promise of utilizing phage cocktails against *Psa*, it highlights the need for rational cocktail design, since the five phages used by Flores *et al*. were highly similar. Their work did not delve into mechanisms of phage resistance developed by *Psa*, but other research towards the development of a phage cocktail for *Pseudomonas aeruginosa* has shown the importance of careful selection of diverse phages for inclusion in cocktails (Yang *et al*., 2020). In particular, the initial interaction of the phage with bacterial surface receptors must be considered, as this will play a key role in the likelihood of bacterial resistance development and its downstream consequences. This was clearly illustrated for *P. aeruginosa* phage PaP1, where phage predation resulted in mutations in the PA1 *wzy* gene, causing O-antigen truncation and phage resistance (Li *et al*., 2018). The consequence of O-antigen truncation in PaP1-resistant PA1 was reduced biofilm formation, thereby decreasing bacterial pathogenicity. Therefore, detailed determination of phage-host interactions enables rational design of phage cocktails that target multiple receptors, which not only attenuate phage resistance development, but also potentially impede bacterial pathogenicity if resistance arises.

We have consistently isolated members of one family of *Psa* phages, the ΦPsa374-like phages, which indicates they are highly abundant in Aotearoa New Zealand (Frampton *et al*., 2014). These phages display biocontrol potential, but despite preliminary analyses of their host range and sequence, little is known about which bacterial receptor is used, the phage receptor binding proteins and the evolution of phage resistance. To understand these important aspects of phage biology for biocontrol design, here we have explored the genomic organisation, virion structure and proteins, host-receptor interactions and both bacterial and phage evolution in response to their interactions. Exploring these phage-host interactions has informed a future path required to exploit these phages in the biocontrol of *Psa* to combat kiwifruit canker that will require partnering these phages with phages utilizing receptors alternative to LPS.

## Results

### Genome sequence of the ΦPsa374 virulent phage

In 2012, multiple phages infecting *Psa* were isolated as potential biocontrol agents (Frampton *et al*., 2014). Among them, phage ΦPsa374, a myovirus isolated from a soil sample in New Zealand, was highly virulent against the plant pathogen *in vitro*. In addition, lysogeny and transduction were not detected previously (Frampton *et al*., 2014), indicating this is a virulent phage and a desirable biocontrol candidate. To further characterize ΦPsa374 we reassembled the phage genome, resulting in a linear dsDNA genome of 98,287 bp with 526 bp direct terminal repeats (DTR), and a G+C content of 47.7%, which was lower than *Psa* (58.4%) (Figure 1A). Genome annotation revealed that ΦPsa374 has 170 predicted open-reading frames (ORFs) and 11 tRNAs (Table S1). The majority of the predicted genes encode hypothetical proteins (128 genes, ~75%). Genes involved in nucleotide metabolism (ribose phosphate pyrophosphokinase *gp29*, dCMP deaminase *gp42*, thymidylate synthase *gp124*, and ribonucleotide reductases *gp126* and *gp127*) and DNA replication (DNA primase *gp105*, DNA polymerase *gp106*, exonuclease *gp115* and endonucleases *gp117*) were predicted. A putative RNA polymerase (*gp64*) was identified, which is also present in other distantly related phages (Garbe *et al*., 2011; Lu *et al*., 2013; Cui *et al*., 2016; Pan *et al*., 2016). In addition, 13 putative structural genes were clustered together with genes involved in packaging and cell lysis (*gp53-84*). These genes are organized in functional modules covering nucleotide metabolism, DNA replication, virion assembly and cell lysis, and the tRNA cluster, in addition to hypothetical proteins (Figure 1A).

**Figure 1.**
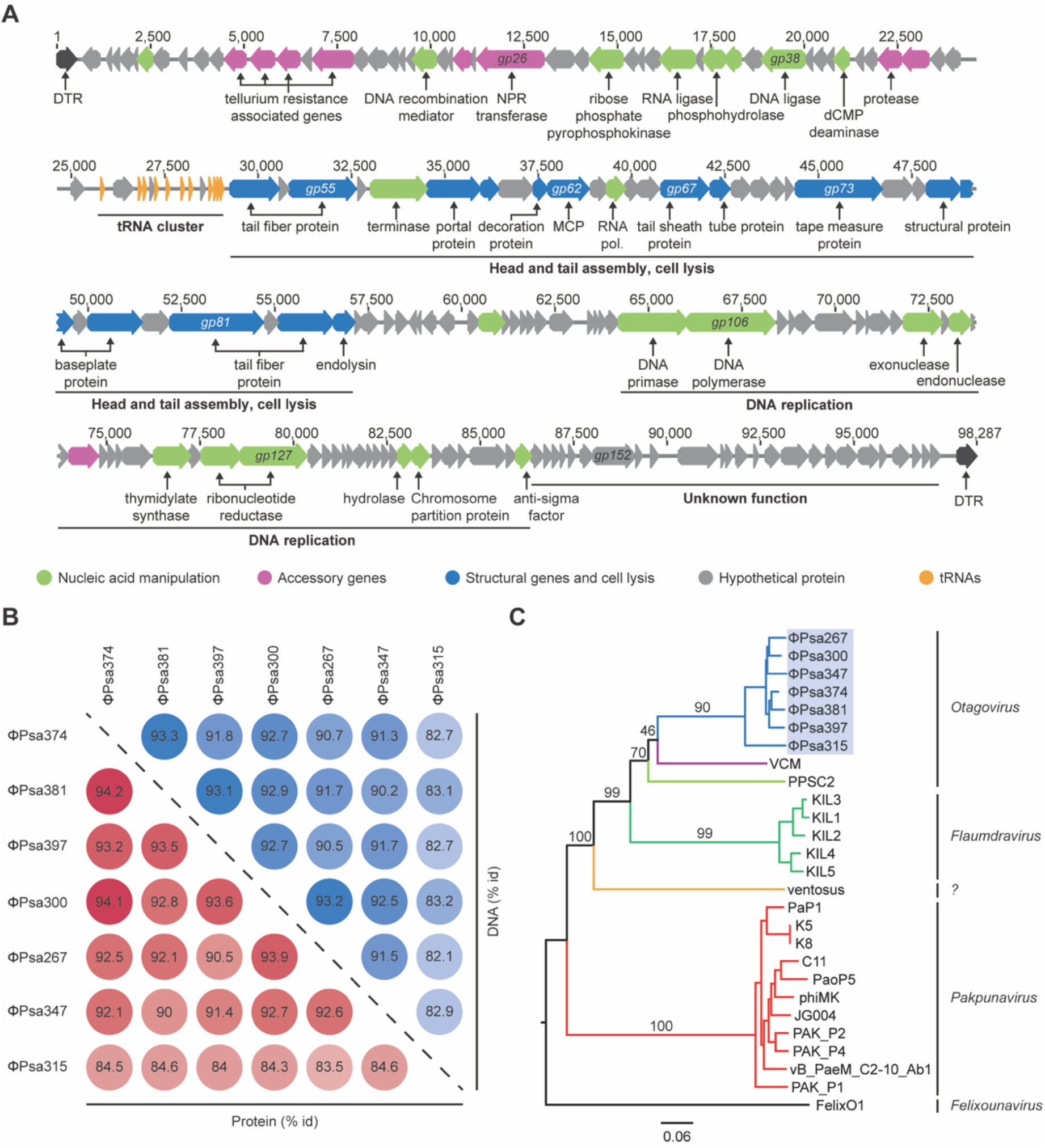
ΦPsa374-like phages are closely related. **A.** Genetic features were classified by function: nucleic acid manipulation (green), accessory genes (pink), structural genes (blue), hypothetical proteins (grey), tRNAs (orange), direct terminal repeats (DTR, black). Function of genes *gp51, gp52, gp67* and *gp68* was predicted from the structural analysis (see below). **B.** Percentage of identical residues at base pairs (blue) and amino acid (red) level between ΦPsa374-like phages. **C.** Genome BLAST Distance Phylogeny tree for ΦPsa374 and related phages. Branch length are scaled according to the distance formula *d_6_*. Branches are color-coded by host: *Pseudomonas* (*P*.) *syringae* pv. *actinidiae* (blue), *P. syringae* DC3000 (violet), *P. fluorescens* (light green), *P. syringae* pv. *porri* (green), *P. syringae* pv. *avii* (orange), *P. aeruginosa* (red), *Salmonella* (black). Phages were grouped into genera according to the International Committee on Taxonomy of Viruses classification. The (?) in the phylogenetic tree represents an unknown genus.

ΦPsa374 also possesses genes associated with tellurium resistance: (*terC, gp11; terD, gp12; terF, gp13* and *telA, gp15*), a rare element reported to have antimicrobial properties (O’Gara *et al*., 1997; Whelan *et al*., 1997; Taylor, 1999; Taylor *et al*., 2002; Hua, 2016). The *ter* genes have also been reported to be involved in colicin and phage resistance (Whelan *et al*., 1995), and are widespread across bacterial phyla (Muñoz-Villagrán *et al*., 2018) and phages, raising questions about their possible role(s) (Moreno Switt *et al*., 2013; Grose *et al*., 2014; Sazinas *et al*., 2016). In ΦPsa374, the *ter* genes do not comprise a complete operon and their function during phage infection are unknown, but are unlikely to provide tellurium resistance due to the lack of all necessary components.

### ΦPsa374-like phages are a discrete group distantly related to other *Pseudomonas* phages

Alongside ΦPsa374, six other ΦPsa374-like phages were isolated from different soil and compost samples in Dunedin, New Zealand (ΦPsa267, 315, 347, 300, 381, 397) (Frampton *et al*., 2014). The phage genomes were assembled *de novo* and a whole genome comparison revealed that the ΦPsa374-like phages are closely related (>82% identity at a DNA sequence) (Figure 1B). While the phages show a conserved genome organization with no major rearrangement, loss and gain of short hypothetical genes are observed across the genomes (Figure S1A). In particular, the tRNA cluster has higher genetic plasticity, with tRNAs gained and lost (Figure S1B & Table S2). The tRNAs encoded by ΦPsa374-like phages have no sequence similarity with bacterial tRNA encoded in *Pseudomonas* strains, but share identity with tRNAs in distantly related *Pseudomonas* phages (Table S2). Phages have been proposed to encode tRNAs to ensure optimal codon usage during infection, and recently work indicates that acquisition of tRNAs by phages is necessary to sustain translation as the host machinery is degraded (Yang *et al*., 2021).

To study the evolutionary relatedness of the ΦPsa374-like phages and other phages, we built a phylogenomic tree based on the Genome-BLAST Distance Phylogeny method. The ΦPsa374-like phages are closely related to other phages from the *Otagovirus* genus that infect *P. syringae* DC3000 (VCM, GenBank ID: LN887844) and *Pseudomonas fluorescens* SA1 (PPSC2) (Wu *et al*., 2018). Other more distant relatives belong to the *Flaumdravirus* (KIL1-like) (Rombouts *et al*., 2016) and *Pakpunavirus* genera (Shen *et al*., 2016) and infect *P. syringae* pv. *porri* and *P. aeruginosa* respectively (Figure 1C). The ΦPsa374-like phages are also related to the unclassified phage ventosus that infects *P. syringae* pv. *avii* (accession number MG018930.1) (Figure 1C). In summary, ΦPsa374-like phages form a highly conserved group within the *Otagovirus* genus, distantly related to other genera of *Pseudomonas* phages.

### Mature ΦPsa374 virions contain 25 structural proteins

To determine the composition of the mature phages, including potential receptor-binding proteins (RBPs), gel-based shotgun proteomic analysis was performed on purified ΦPsa374 virions. Peptide sequences detected by LC-MS/MS analysis were mapped back to the translated genes of ΦPsa374 (Figure 2A), revealing that 22 out of 25 hits are proteins encoded in the predicted structural locus (Figure 2B and Table 1). The major capsid protein (Gp062) was the most abundant, followed by a structural protein (Gp067), and a hypothetical protein (Gp068). Their close proximity to the tape measure protein (Gp073) and relative abundance suggest that these proteins are involved in tail or sheath formation. In addition, Gp061, a homologue of the lambda head decoration protein D, and the two tail fibers Gp083 and Gp081 (predicted RBPs) had the 4^th^, 5^th^ and 6^th^ highest relative abundance, respectively. Proteomics of the distantly related *P. aeruginosa* phages PaoP5 (Shen *et al*., 2016), PAK-P3 (Henry *et al*., 2015) and PaP1 (Lu *et al*., 2013) and the *P. syringae* pv. *porri* KIL-phages identified multiple analogous proteins of similar size (Rombouts *et al*., 2016) (Table S3).

**Figure 2.**
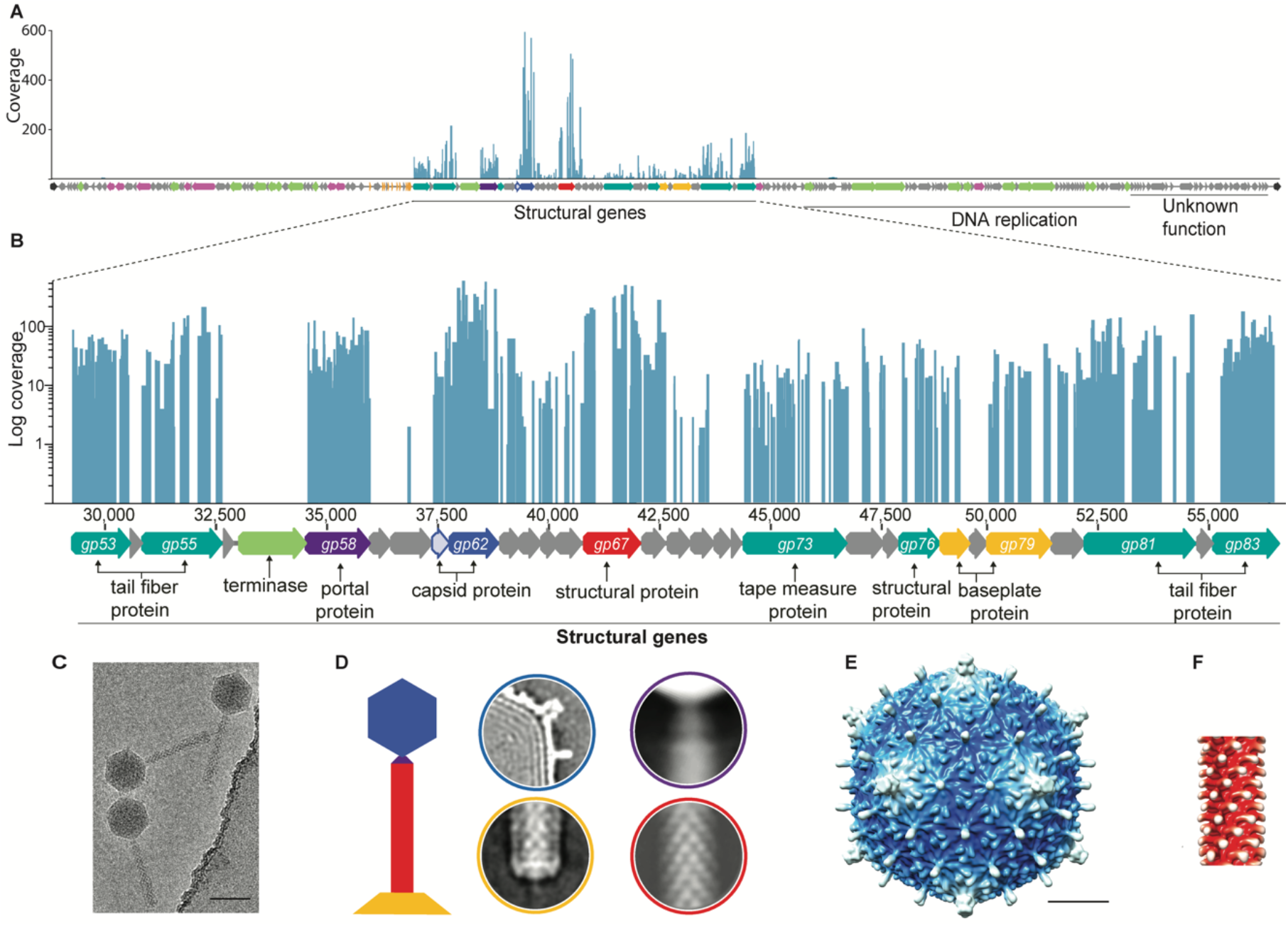
Structural analysis and cryo-EM of ΦPsa374 viral particles. **A.** Peptide sequences from mature virions mapped back to the genome of ΦPsa374 and **B.** log-fold coverage of peptide sequences mapped to the structural region of the genome. **C.** A region from a cryo-EM micrograph showing ΦPsa374 particles, scale bar is 750 Å. **D.** A schematic representation of a bacteriophage architecture, the main modules identified in the cryo-EM are shown in the same colors with the capsid (blue), neck (violet), tail (red), and baseplate (yellow). The local average shows that the ΦPsa374 baseplate has a dimension similar to the of the tail. Due to their flexibility, tail fibers are not visible in 2D averages. **E.** Two dimensional average images showing the structural isosurface representations of ΦPsa374 capsid and **F.** a segment of the tail. Scale bar is 200 Å.

**Table 1.** Structural proteome of ΦPsa374 by LC-MS/MS of isolated phage virions.

| Locus tag | Description | Closest *Pseudomonas* Phage Homologue |  | % identity (amino acid) | Domains or PHA#<sup>1</sup> | Unique Peptides<sup>2</sup> | AUC<sup>3</sup> | MW<sup>4</sup> (kDa) | Coverage (%) |
| --- | --- | --- | --- | --- | --- | --- | --- | --- | --- |
|  |  | Phage | Locus tag |  |  |  |  |  |  |
| gp053 | SGNH-hydrolase (Tail Fiber protein) | VCM | VCM_00116 | 84.24 | pfam13472 | 42 | 1.51E+08 | 44.7 | 94 |
| gp055 | Tail fiber protein | - | - | - |  | 44 | 1.40E+08 | 63 | 74 |
| gp058 | Portal protein | VCM | VCM_00113 | 92.84 |  | 72 | 7.77E+07 | 54.5 | 95 |
| gp061 | Phage lambda head decoration protein D | VCM | VCM_00110 | 91.41 | pfam02924 | 23 | 3.49E+08 | 13.7 | 91 |
| gp062 | Major capsid protein | KIL4 | FDI83_gp156 | 90.65 | pfam03864 | 124 | 5.53E+09 | 39.5 | 99 |
| gp063 | Phage protein | VCM | VCM_00108 | 86.54 |  | 11 | 1.93E+07 | 18 | 78 |
| gp064 | Hypothetical protein | VCM | VCM_00107 | 77.42 |  | 4 | 6.40E+05 | 17.1 | 39 |
| gp065 | Hypothetical protein | VCM | VCM_00106 | 95.9 |  | 12 | 5.72E+06 | 13.9 | 79 |
| gp066 | Hypothetical protein | VCM | VCM_00105 | 84.13 |  | 5 | 2.11E+07 | 21.7 | 38 |
| gp067 | Structural protein | VCM | VCM_00104 | 82.44 | pfam11863 | 103 | 1.89E+09 | 45.8 | 76 |
| gp068 | Hypothetical protein | VCM | VCM_00103 | 92.49 | pfam11681 | 26 | 7.57E+08 | 18.7 | 95 |
| gp069 | Hypothetical protein | KIL1 | vB_PsyM_KIL1_0064 | 88.02 |  | 4 | 4.56E+06 | 18 | 19 |
| gp070 | Hypothetical protein | PPSC2 | PPSC2_15 | 78.53 |  | 5 | 1.56E+07 | 19 | 44 |
| gp073 | Tape measure protein | PPSC2 | PPSC2_18 | 59.67 | HI1514 | 53 | 3.37E+07 | 84.4 | 64 |
| gp074 | Hypothetical protein | VCM | VCM_00098 | 63.34 |  | 18 | 4.24E+07 | 29.4 | 33 |
| gp076 | Structural protein | KIL1 | BH774_gp130 | 85.9 |  | 17 | 2.76E+07 | 34.1 | 60 |
| gp077 | Baseplate protein | KIL5 | vB_PsyM_KIL5_0079 | 79.05 | pfam06890 | 10 | 1.19E+07 | 27.4 | 36 |
| gp078 | Tail lysozyme | VCM | VCM_00094 | 78.51 |  | 2 | 3.07E+06 | 13.4 | 21 |
| gp079 | Baseplate J-like protein | VCM | VCM_00093 | 81.16 | pfam04865 | 25 | 5.80E+07 | 51.7 | 61 |
| gp080 | Phage protein | PPSC2 | PPSC2_25 | 89.71 | pfam11041 | 22 | 4.96E+07 | 26.6 | 83 |
| gp081 | Tail fiber protein | PPSC2 | PPSC2_26 | 62.27 |  | 60 | 1.82E+08 | 85.5 | 66 |
| gp083 | Tail fiber protein | VCM | VCM_00089 | 70.85 | PHA02584 | 51 | 2.92E+08 | 51.7 | 82 |

<sup>1</sup>Protein domains classified into families (pfam). <sup>2</sup>Unique peptides identified per protein. <sup>3</sup>The area under the curve (AUC) represents the relative protein abundance measured by the TOP3 approach. <sup>4</sup>MW is the molecular weight of the protein.

The genes encoding the structural proteins for ΦPsa374 and the ΦPsa374-like phages showed high levels of synteny with related *Pseudomonas* phages (Figure S1A&C). ΦPsa374-like phages contained a SGNH-hydrolase, which was not present in any of the related phages. In addition, the KIL phage family and JG004 possessed either one or more endonucleases that were absent in the other phages. All differences in genes encoding the structural proteins for ΦPsa374 and related *Pseudomonas* phages are summarised in Table S3. Identification of structural proteins in ΦPsa374 virions and correlation with orthologous proteins in genetically similar phages helps to determine which proteins are potentially involved in binding to phage receptors on the bacterial cell surface.

### The ΦPsa374 capsid possesses a T=9 triangulation

To understand the overall bacteriophage architecture we performed cryo-electron microscopy (cryo-EM) on ΦPsa374 particles (Figure 2C&D). The 3D reconstruction of the capsid (11.5 Å resolution) showed an icosahedral arrangement with a maximum diameter measured between two five-fold axis of 800 Å (Figure 2E). The capsid displays a triangulation number of T=9, uncommon in the *Myoviridae* family. This capsid geometry was previously reported in *Shigella* myophages (Doore *et al*., 2018), in coliphage N4 (*Podoviridae*) (Choi *et al*., 2008), *Bacillus* phage Basilisk (Grose *et al*., 2014) and *Mycobacterium* phage Rosebush (Podgorski *et al*., 2020) (both members of the *Siphoviridae* family). This larger triangulation number facilitates the packing of a larger genome of ~98 KB (Doore *et al*., 2018; Kamiya *et al*., 2021). At the interior of the capsid the arrangement of the dsDNA genome is visible as a series of layers separated by ~23 Å (Figure 2D). Based on sequence similarity and the abundance data derived from mass spectrometry, Gp062 constitutes the major capsid protein (MCP). A sequence alignment indicates (data not shown) that Gp062 contains the usual HK97 fold conserved among the MCP of dsDNA phages (Wikoff, 2000; Duda and Teschke, 2019). Hexamers formed by ΦPsa374 MCPs have an average thickness of ~40 Å and display an elongated decoration protein at their centre (Figure 2E). Based on molecular weight and relative abundance, we propose that Gp061 is a decoration protein, as its shape is similar to decoration proteins reported for the *Pectobacterium* phage ΦTE (Watson *et al*., 2019) and coliphage T5 (Vernhes *et al*., 2017).

ΦPsa374 particles have a tail with a total length of ~1300 Å that connects to the capsid through a thin neck at one of the five-fold axes (Figure 2D). The tail tube has a length of 1100 Å and is formed by 30 disks with a thickness of 37 Å and a C6 symmetry, giving the tail a six-start helical symmetry with a twist of 25.5° (Figure 2D&F). At the interior of the sheath, the inner tube proteins form a central channel with a diameter of ~40 Å. Considering the stoichiometry indicated by our structural analysis, the relative abundance from the mass spectrometry data, and the sequence alignments, we propose that Gp067 is the sheath protein, while Gp068 is the tube protein, confirming mass spectral data. Lastly, at the distal end, the base plate region shows few features, displaying a diameter similar with that of the tube (Figure 2D). The structure of the tail fibers was not possible to determine by cryo-EM due to their flexibility, which impeded their visualisation in 2D averages. In summary, here we identified that ΦPsa374 particles are comprised of 25 structural proteins and display an uncommon T=9 triangulation.

### ΦPsa374-like phages employ LPS as a receptor *in vitro* and *in planta*

Recognition and adsorption to bacterial receptors is a critical step in phage infection (Bertozzi Silva *et al*., 2016; Hampton *et al*., 2020). To identify the receptor of ΦPsa374-like phages we tested their ability to infect *Psa* transposon (Tn) mutants with insertions in genes in the LPS and flagella biosynthetic pathways (Mesarich *et al*., 2017). LPS and flagella are both common bacterial cell surface molecules exploited by phages for binding and infection (Stone *et al*., 2019). Most of the transposon mutants had inserts in genes in the common polysaccharide antigen locus (Jayaraman *et al*., 2020), which encodes proteins for the biosynthesis and transport of the O-antigen portion of LPS across the outer membrane (Figure 3A). All ΦPsa374-like phages had major reductions in their ability to infect the LPS mutants, but plaqued efficiently on the flagellar P-ring protein mutant (*flgl::Tn*) (Fig 3B). In contrast, a control flagella-tropic *Psa* phage ΦPsa21 (Frampton *et al*., 2014; Wojtus *et al*., 2019) infected all LPS mutants but not the flagella mutant (Fig 3B). These results indicate than an unidentified component of the *Psa* LPS acts as a major receptor for the ΦPsa374-like phages.

**Figure 3.**
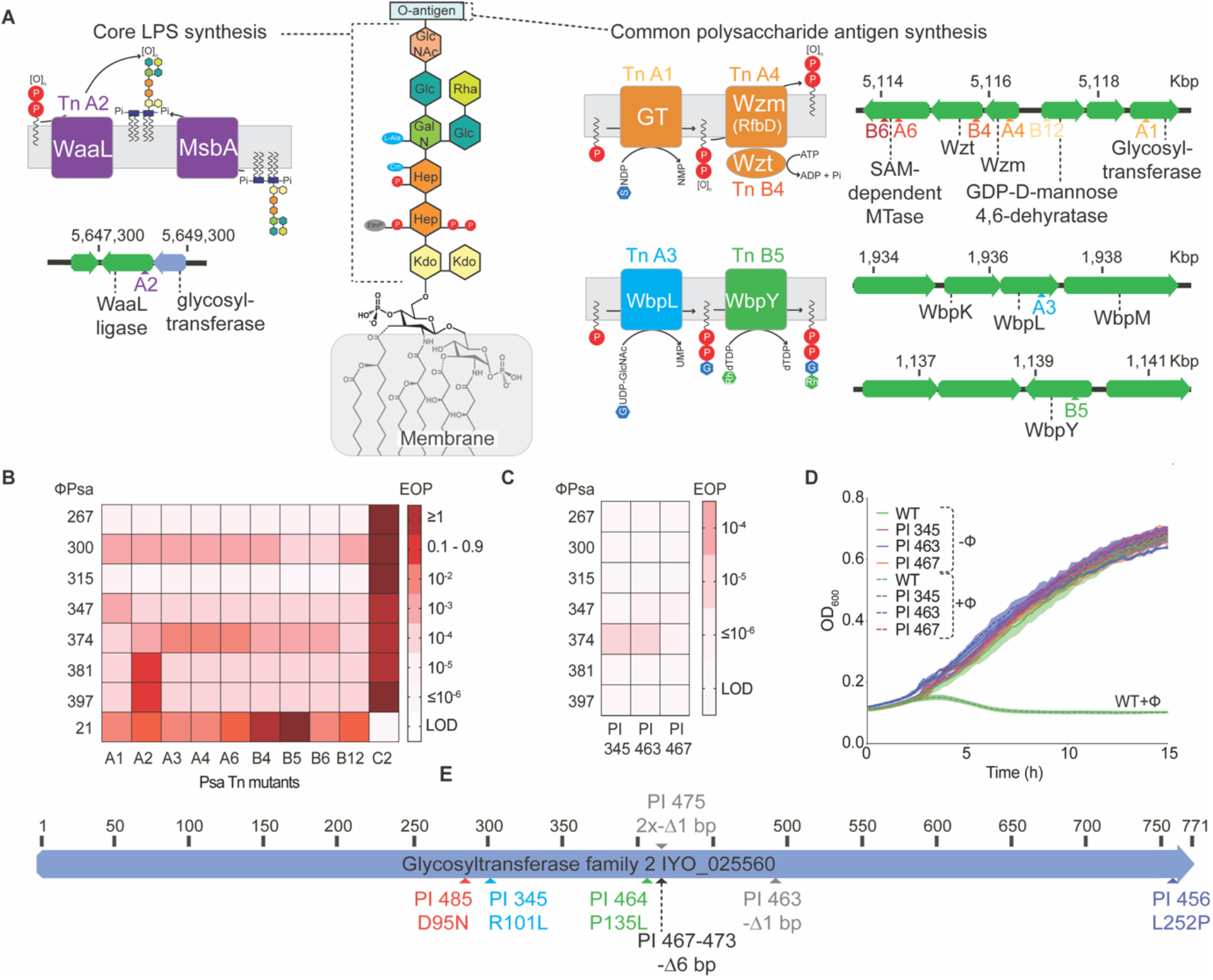
ΦPsa374-like phages use LPS as a receptor *in vitro* and *in planta*. **A.** A schematic representation of the structure of LPS in *P. syringae*, the core polysaccharide is based on structural elucidation work done on *P. syringae* pv. *phaseolicola* (Zdorovenko *et al*., 2004). Schematic representation of Tn insertions in genes in the core LPS locus (purple) and common polysaccharide antigen pathway (orange, blue and green). Tn sites and predicted role of key proteins in their cellular context are shown. Color coding differentiates location of genes within the genome and genes with insertions are shown schematically. For simplicity, Transposon mutants are named e.g. either Tn A1 or A1, interchangeably. The proteins disrupted by A6, B6 or B12 are omitted for simplicity. **B.** Sensitivity of a panel of *Psa* LPS-Tn mutants against ΦPsa374-like phages and ΦPsa21 (a flagella tropic *Psa* jumbo phage included as a control). EOP is efficiency of plaquing. Darker red indicates greater efficacy of a phage (higher EOP) to infect a strain. LOD (limit of detection) indicates no plaques were observed. Strain C2 has a Tn-insertion in a gene encoding the flagella P-ring protein (Flgl) and acts as a control that is sensitive to all phages except ΦPsa21. Sensitivity of *Psa* phage resistant plantlet isolates (PI) from *in planta* studies against the ΦPsa374-like phages in **C.** plate EOP assays and **D.** liquid phage infection assays. In D. a multiplicity of infection (MOI) of 0.01 was used. In B-D data was repeated in biological triplicate, solid or dotted lines are the mean while shading of these represents the standard error of the mean (SEM). Raw EOP data is presented in Figure S4. **E.** Locations of mutations in the glycosyltransferase gene within the core LPS locus in 9 *Psa* isolates sampled from ΦPsa374-treated kiwifruit plantlets (details in Table S4).

To determine the suitability of ΦPsa374 as a biocontrol agent, we assessed the efficacy of the phage to control *Psa* infection in kiwifruit plantlets (Figure S2A). ϕPsa374 was applied on days 1, 2, 4 and 8, while *Psa* was inoculated on day 2, just prior to the second phage application. In plantlets treated only with *Psa*, bacterial numbers increased by ~10^4^ from day 2 to 9, plateauing between days 9 to 11 with a *Psa* density of ~1 x 10^7^ CFU/mm^2^. In contrast, the *Psa* density on ϕPsa374 treated plantlets was significantly delayed and reduced to below the detection limit until day 4 (between untreated and phage treated plantlets). By day 5 *Psa* began to increase on ϕPsa374 treated plantlets. From day 8 to 16 no significant difference in *Psa* density between untreated and phage treated plantlets was observed. Notably, the phage titre remained stable throughout the experiment at ~10^7^ pfu/mm^2^ (Figure S2A). We hypothesized the *Psa* isolates emerging from day 5 onwards were ϕPsa374-resistant. This delayed *Psa* growth shows that ΦPsa374 possesses potential as a biocontrol agent, but when employed alone, we predict that bacterial resistance can arise.

To determine if the Psa isolates that survived ΦPsa374 treatment *in planta* were due to phage resistance, we isolated *Psa* from both the phage treated and untreated plantlets. Untreated *Psa* remained sensitive to ϕPsa374 infection (Figure S2B&C). In contrast, ΦPsa374-treated isolates showed the emergence of ΦPsa374-resistance from day 4. From day 5 onwards all isolates screened were ΦPsa374-resistant (Figure S2B&C). From this pool of ΦPsa374-resistant isolates, 3 representatives from day 11 and 16, were further characterized. Plaquing assays showed high levels of resistance to all ΦPsa374-like phages, further corroborated by liquid infection assays which showed resistance to ΦPsa374 (Figure 3C&D). To determine whether LPS mutations were the cause of phage resistance *in planta*, we sequenced the whole genomes of nine plant derived isolates (PI), details of which can be found in Table S4. The nine isolates had seven distinct mutations in a single gene (locus tag IYO_025560) (Templeton *et al*., 2015) (Figure 3E and Table S4). This protein belongs to the glycosyltransferase (GT-A) superfamily, responsible for catalysis of glycosidic bonds in oligosaccharides, polysaccharides and glycoconjugates (Schmid *et al*., 2016). GT-A enzymes, including IYO_025560, possess an α/β/α sandwich fold, and commonly exhibit a DxD motif (Figure S3), which uses a manganese cofactor to bind sugar-nucleoside phosphates for the addition of sugars to sugars, phosphates or proteins (Wiggins and Munro, 1998; Breton *et al*., 2006). The location of this gene within an operon immediately upstream of the *waaL* O-antigen ligase (Jayaraman *et al*., 2020) (Figure 3B) implies that it forms glycosidic bonds between terminal sugars of the core LPS or between the core terminal sugars and the O-antigen. Some bacterial isolates from the *in planta* experiments had single-nucleotide polymorphisms (SNPs) that caused non-synonymous mutations, others had small deletions that caused a frameshift resulting in truncated proteins and finally, three identical isolates (from the same plantlet) harboured a 6 bp deletion causing the deletion of tyrosine and valine residues (Figure 3E and Table S4). Notably, the GT-A gene is within the same operon as the *waaL* gene, which when mutated (Tn A2 mutant) also led to phage resistance (Figure 3B). Therefore, loss of function or mutations in these genes result in resistance against ΦPsa374-like phages. It is likely that resistance is manifested through alterations in the structure or abundance in the LPS of these mutants.

### ΦPsa374-resistant *Psa* have altered LPS but virulence is not impaired

Previously, the *Psa* mutants with transposon insertions in LPS biosynthetic pathway genes were shown to lack LPS (Hockett *et al*., 2017; Jayaraman *et al*., 2020). To determine whether this was also the case for the ΦPsa374-resistant plantlet isolates, we compared their LPS profiles to those of both wild-type (WT), five LPS transposon mutants (Figure 4A), and an untreated plantlet isolate (PI 406) which remained sensitive to ΦPsa374 (Figure 4A). Both the WT and PI 406 (Lanes 1, 12 and 7 respectively) showed the characteristic laddering for intact LPS. In contrast, most of the transposon mutants lacked LPS irrespective of the insert location (common and core LPS biosynthetic pathways) (Figure 3A). An exception was the A2 mutant, which had reduced LPS levels. This strain has a mutation in *waaL* that encodes a protein involved in synthesis of the core domain. The *Psa* ΦPsa374 resistant plantlet isolates (PI 345, 463, 475 and 485) expressed less LPS than the WT and the LPS bands ran with a slightly different electrophoretic mobility (Figure 4A). This implies that mutations in a single GT-A due to ΦPsa374-resistance result in an alteration of chemical composition and thinning of LPS in *Psa* in response to phage predation.

**Figure 4.**
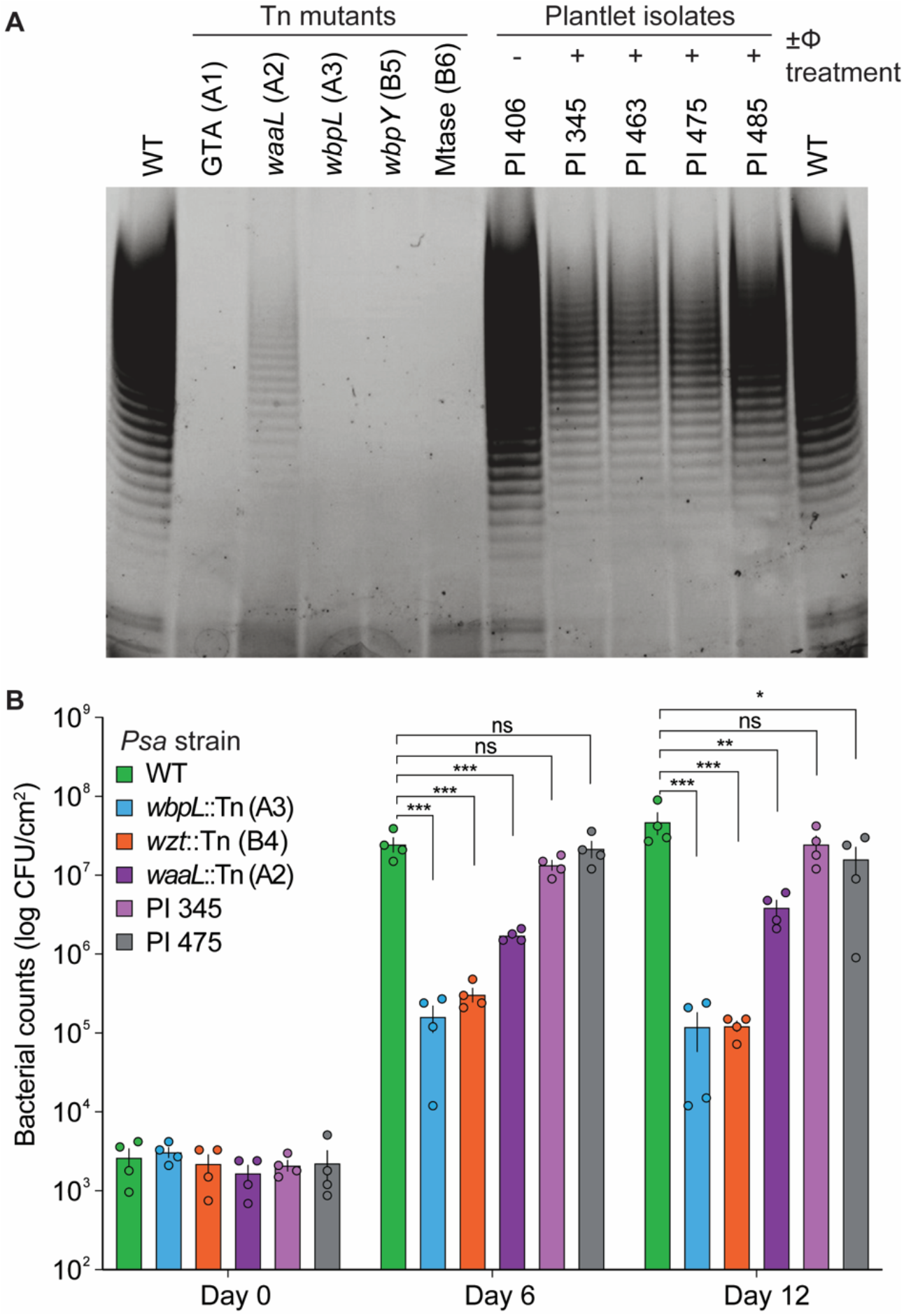
ΦPsa374-resistant *Psa* have different LPS and pathogenicity profiles from both wildtype and LPS Tn5-mutants. **A.** Lipopolysaccharide (LPS) was resolved using a 4%–2% premade NuPAGE™ Bis-Tris SDS gel and stained with Pro-Q™ Emerald 300 Lipopolysaccharide Gel Kit. Lane 1, WT (*Psa* ICMP 18884); lane 2, A1, (IYO_23030::Tn, glycosyl-transferase (GTA) involved in Drhamnose biosynthesis, common LPS); Lane 3, A2 (IYO_025555::Tn, core LPS); Lane 4, A3 (*wbpL::Tn*, common LPS); Lane 5, B5 (IYO_005330::Tn, common LPS); lane 6, B6 (IYO_023005::Tn, SAM-dependent methyl-transferase (Mtase), common LPS); lane 7, PI 406; Lane 8, PI 345; Lane 9, PI 463; Lane 10, PI 475; Lane 11, PI 485; Lane 12, WT (*Psa* ICMP 18884). **B.** *In planta* bacterial growth on *Actinidia chinensis* ‘Hort16A’ was recorded at days 0, 6 and 12. The identity of the *Psa* isolates is shown in the key. Each data point represents a biological replicate, n=4. One-way ANOVA multiple comparisons were performed on data, ns (not significant) = adjusted P value > 0.05, * = adjusted P value 0.05 – 0.005, ** = adjusted P value 0.005 – 0.001, *** = adjusted P value < 0.001.

Next, we tested a set of both transposon and ΦPsa374-resistant plantlet isolates *in planta* for their pathogenicity on kiwifruit (Figure 4B). Growth of the *Psa* strains *in planta* showed that null LPS mutations were reduced >100-fold compared with the WT. In contrast, the spontaneous ΦPsa374-resistant plantlet isolates retained their capacity to grow *in planta* with only slightly lower growth than the WT. Bacterial density presented here qualitatively correlated with pathogenicity, as leaf lesions associated with *Psa* infection were more abundant in plantlets infected with WT and plantlet isolates (Figure S5). In contrast, no significant disease was noted in plantlets infected with LPS-Tn mutants of *Psa*. Interestingly, the A2 mutation in the gene encoding *waaL* had an intermediate effect on the ability of *Psa* to grow on kiwifruit. This data shows that the development of PsaΦ374-resistance *in planta* results in genotypic and phenotypic differences in LPS in *Psa* and that these phage resistant isolates can still cause significant disease in kiwifruit plantlets.

### *In vitro* evolution of ΦPsa374 with phage-resistant bacteria overcomes receptor resistance

For phages to overcome bacteria that have mutated phage receptors, they can adapt in a coevolutionary dynamic through mutations to their structural proteins, including receptor binding proteins (RBPs) (Burmeister *et al*., 2016; Yuan *et al*., 2019). To investigate how ΦPsa374 interacts with *Psa* and whether it can evolve to overcome the LPS mutants, we performed *in vitro* evolution experiments. ΦPsa374 was propagated on the WT strain and used to infect the bacterial plantlet isolate PI 345 (Figure 5A). While PI 345 was resistant to ΦPsa374, plaques produced by ΦPsa374 mutants (termed escape phages (escΦ)) were detected. A single escΦ plaque was propagated on PI 345 three times to ensure purity. The escΦ was able to equally infect both WT *Psa* and the bacterial plantlet isolate PI 345 on plates or in liquid culture (Figure 5A&B). Importantly, passaging the escΦ back through WT *Psa* and then using this to infect both WT and the plantlet isolate showed that the escΦ still infected both to an equivalent level. This indicates that the escΦ is due to genetic differences relative to the original ΦPsa374 and is not a result of phenotypic escape.

**Figure 5.**
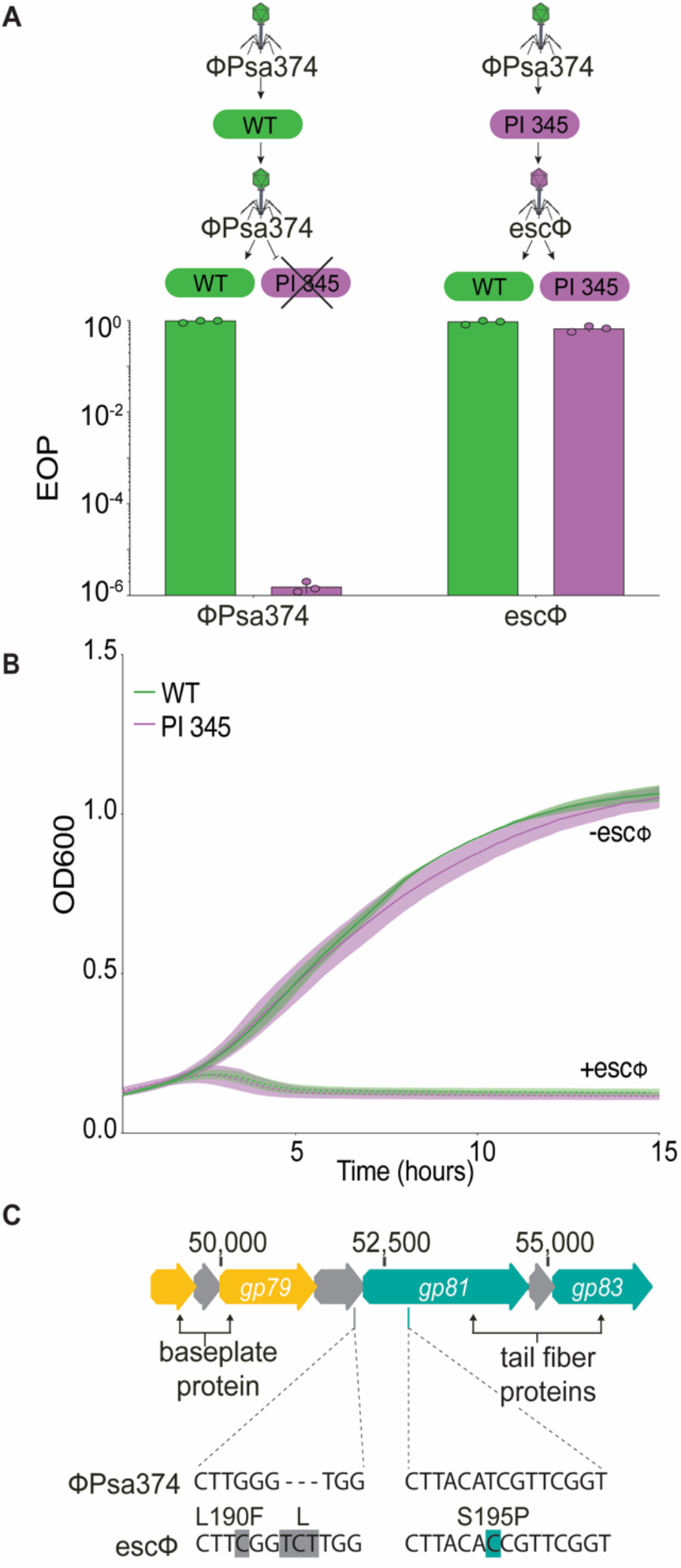
An escape phage/ΦPsa374-mutant overcomes receptor-mediated phage resistance by mutations in a tail fiber and uncharacterized structural protein. **A.** EOP data for ΦPsa374 and an escape phage (escΦ) against PI 345. Each bar represents the mean of biological triplicates, with each individual data point plotted to show variation. **B.** Growth curve data for the escΦ against WT (ICMP 18884) and PI 345, showing restoration of infection in PI 345. Data was measured in biological triplicates, solid or dotted lines are plotted as the mean while shading represents the SEM of the mean. **C.** Location of the escΦ mutations in two structural genes, *gp080* and *gp081*.

Whole genome sequencing of the escΦ revealed non-synonymous SNPs in three genes (*gp039* (D18Y), *gp080* (L190F) and *gp081* (S195P)) (Figure 5C). In addition to a SNP, *gp080* has a 3 bp insertion, resulting in a leucine insertion at amino acid position 192. In the attachment stage, structural proteins interact with phage receptors on the bacterial cell surface. Therefore, it is expected that genetic phage escapes would have mutations in structural genes. Since Gp039 was not identified by proteomics of the mature virions (Figure 2), it is unlikely to be involved in the escape phenotype. On the other hand, Gp080 and Gp081 were identified in mass spectral data, indicating they are part of the phage particle (Figure 2B). The presence of Gp080 and Gp081 in virions, and their observed mutations in the escape mutant phage, implies that these proteins play a role in host recognition of *Psa* by ΦPsa374.

To further characterize and validate this escape phage genotype, we reinfected PI 345 with the escΦ and isolated two plaques. We then confirmed mutations in *gp080* and *gp081* by PCR and Sanger sequencing (Figure S6), which revealed both plaques have identical mutations as seen in WGS (Figure 5C). After propagating these phages through PI 345, we tested their ability to infect the flagellum Tn-mutant (C2) and three selected plantlet isolates (Figure S4B). Similar to results presented for ΦPsa374 and ΦPsa374-like phages (Figure 3B and C), these escape phage replicates (rep 1 and 2) were capable of infecting the flagellum mutant. Both escape phage replicates infected the ΦPsa374-resistant PI 345 and 467 isolates, but no plaques were detected on PI 463. PI 345 and 467 show a SNP and insertion in the GT-A (IYO_025560), respectively, while PI 463 has a single base pair deletion which results in a truncation of the glycosyltransferase (Figure S3). LPS profiles for these plantlet isolates are indistinguishable in Figure 4A; therefore, further work is required to elucidate precise structural differences in LPS between the isolates that influence phage infectivity. Our data demonstrates that subtle mutations in the host GT-A can be overcome by minor mutations in RBPs of the phage.

## Discussion

Here we report a detailed analysis of the ΦPsa374-family of potential biocontrol phages that infect *Pseudomonas syringae* pv. *actinidiae* (*Psa*), an important phytopathogen of kiwifruit. Comparative genomics revealed that these phages form a discrete *Otagovirus* genus within the *Felixounavirinae* phages, which is related to other *Pseudomonas* phage genera (*Flaumdravirus* and *Pakpunavirus*). The ΦPsa374 structural proteome was similar to other *Felixounavirinae Pseudomonas* phages (Lu *et al*., 2013; Henry *et al*., 2015; Rombouts *et al*., 2016; Shen *et al*., 2016). Cryo-EM revealed that the phage capsid has icosahedral symmetry with two hexavalent capsomers between two adjacent five-fold axes, indicating a T=9 triangulation number. This uncommon T number has only been reported in a few other phages (Choi *et al*., 2008; Grose *et al*., 2014; Doore *et al*., 2018; Podgorski *et al*., 2020). The ΦPsa374-family of phages utilize LPS as a major receptor and mutations within this biosynthetic pathway can emerge during *in planta* infection assays with ΦPsa374 phage treatment. In response, ΦPsa374 can evolve RBP mutations that enable phage resistance to be overcome. This work provides insight into the bacterial response to ΦPsa374 that will help to inform the rational design of phage cocktails for *Psa* biocontrol that utilize different receptors on *Psa*. The apparent natural abundance and frequent isolation of the ΦPsa374-like phages in New Zealand soil and leaf samples implies stability in environmental conditions similar to kiwifruit orchards.

The exploitation of LPS as a receptor by ΦPsa374-like phages occurs in other *Felixounavirinae Pseudomonas* phages, including PaP1 (Li *et al*., 2018), JG004 (Garbe *et al*., 2011) and the KIL-family (Holtappels *et al*., 2020), and tailocins (Carim *et al*., 2021). By sequencing ΦPsa374-resistant *Psa* isolates that arose *in planta*, we identified mutations in a GT-A glycosyltransferase in the core LPS locus. Typically, mutations in multiple genes involved in LPS biosynthesis can lead to phage resistance (Meaden *et al*., 2015; Li *et al*., 2018; Olszak *et al*., 2019; Holtappels *et al*., 2020), similar to our *in vitro* resistance data with the transposon mutants. It was therefore surprising that multiple independent mutations were only found in a single GT-A glycosyltransferase in the core LPS biosynthetic loci when phage-resistant bacterial isolates from the *in planta* experiment were sequenced. LPS from these isolates was less abundant and had altered electrophoretic mobilities, suggesting a change in chemical composition. A minor phenotypic mutation such as this could explain the apparent loss of recognition of LPS by the phage tail fibers, while still allowing these strains to maintain pathogenic growth on kiwifruit similar to that observed in WT. In contrast, the LPS transposon mutants were significantly impaired for growth and virulence. The development of phage-resistance in *Psa* while maintaining pathogenicity is an important finding of this work. It means that ΦPsa374-like phages need to be used in a cocktail with other phages that target different receptors to reduce resistance or bacterial virulence. This is in direct contrast to the work of Hernandez *et al*. (Hernandez and Koskella, 2019), which showed the development of phage resistance impaired pathogenicity and the ability of mutants to out-compete WT bacteria *in planta*.

Our data suggests two possible scenarios upon cessation of ΦPsa374-treatment. First, any remaining rare WT *Psa* may eventually outcompete the phage-resistant mutants. This would result in a *Psa* infection re-treatable with ΦPsa374, but which would likely again result in phage resistance. Second, the spontaneous phage resistant strains, which dominate following ΦPsa374-treatment, might compete effectively with the WT and cause disease. Ultimately, both scenarios would require further phage treatment with phages that infect these emergent phage resistant strains and hence, the use of cocktails or evolved escape ΦPsa374 phages is needed. Below, we discuss the possibility of isolating phages for the latter approach.

ΦPsa374 was able to overcome phage resistance mediated by mutations in LPS by acquiring mutations in *gp080* and *gp081*. A SNP and insertion were found towards the C-terminus of Gp080, while the SNP in Gp081 is located at the N-terminus. Phage tail fiber structures typically show conservation of the N-termini, which act as an attachment point to the phage, and hypervariability or mutations at the C-termini, that act as RBPs (Dams *et al*., 2019; Latka *et al*., 2019; Boon *et al*., 2020). Indeed, this is what is observed for *gp081* for the ΦPsa374-like family (Figure S7). Homologs of Gp081 are annotated as tail fiber RBPs: Gp084 (phage JG004, host PAO1) and Gp069 (phage PaP1, host PA1) (Garbe *et al*., 2011; Le *et al*., 2013)). Le *et al*. determined Gp084_JG004_ and Gp069_PaP1_ acted as RBPs due to variation at the C-terminus of the tail fibers and the capacity to alter the host of PaP1 from PA1 to PAO1 by cloning the C-terminus of *gp084*_JG004_ into *gp069*_PaP1_. The other gene of interest *gp080* has an unknown function and belongs to the DUF2612 family. While the function of *gp080* is unknown, mutations in an orthologous gene (*orf75*) in *Pseudomonas* phage PaoP5, resulted in a genotypic escape from phage-resistant mutants with truncated O-antigen (Yang *et al*., 2020). The location of the mutation in *gp081* implies a change in function of protein rather than alteration of host specificity, and may even be deleterious as proline substitutions often cause location-dependent destabilization or disrupt protein folding (Bajaj *et al*., 2007). In contrast, the SNP and insert in Gp080 at the C-terminus suggests that this protein has been altered in response to changes in LPS. Our data shows that while *Psa* can become resistant to ΦPsa374-like phages, this can be overcome through mutations in structural proteins/tail fibers in ΦPsa374 to produce an alternative binding mode. Our work highlights the need to examine phage-host interactions *in planta* to properly realize evolutionary adaptation of bacteria to phage predation in relevant ecological contexts. This research illustrates the suitability of ΦPsa374-like phages as a biocontrol of *Psa*, and demonstrates the importance for rational design of cocktails that include phages that have been evolved to overcome resistance and/or additional phages that target alternative receptors.

## Experimental Procedures

### Bacterial strains, culture conditions and phage stock preparation

*Pseudomonas syringae* pv. *actinidiae* ICMP18884 (*Psa*) (Templeton *et al*., 2015) (accession number: CP011972, chromosome and CP011973, plasmid) was grown in Lysogeny Broth (LB) at 28°C in shaking conditions (160 RPM), or on solid LB agar (LBA) containing 1.5% w/v agar. Strains used in this study are listed in Table S5 and S6.

Phages were isolated from different soil and compost samples from Dunedin, New Zealand, as previously described (Frampton *et al*., 2014). Phage propagation was performed on LBA using an agar overlay method. Briefly, 10-fold dilutions of a high titre ΦPsa374 stock were plated onto an LBA soft overlay (0.35% w/v agar) previously seeded with *Psa*. The plate was incubated overnight at 28°C and plaque formation was observed to determine which dilution would produce almost confluent lysis. A 100 μL aliquot of a *Psa* overnight culture and 100 μL of the diluted phage stock, were added to 4 mL of molten soft LBA (0.35% w/v) and poured onto LBA plates. The phage stock concentration was enough to produce almost confluent lysis. The plates were incubated overnight at 28°C and the soft agar was scrapped from the plate and pooled into centrifugation tubes. A few drops of chloroform (NaCO_3_ saturated) were added and mixed by vortexing to lyse the cells. Cell debris were removed by centrifugation (2,000 *g* for 20 min at 4°C) and the phage containing supernatant was transferred to a sterile tube where chloroform (NaHCO_3_ saturated) was added to prevent bacterial growth. To titrate the phage lysates, 10-fold dilutions were spotted onto a soft LBA overlay previously seeded with *Psa* and incubated overnight at 28°C. Phage titre was reported as PFU/mL and stocks were stored at 4°C.

### Phage genome assembly and annotation

Previously, phage genomic DNA (gDNA) was extracted and sequenced (Frampton *et al*., 2014). The resulting reads were individually assembled *de novo* into one contig for each ΦPsa374-like phage using Geneious 6.1.6 using default settings, and direct terminal repeats were identified with PhageTerm (Garneau *et al*., 2017). The phages were annotated using RAST and tRNAs were identified with tRNAscan-SE (v2.0) (Lowe and Chan, 2016). Open reading frames (ORFs) were further annotated by manual curation using BLASTN, BLASTP and PSI-BLAST (McGinnis and Madden, 2004) and InterProScan (Quevillon *et al*., 2005). The genomes were visualized using Geneious 11.1.4.

### Nucleotide sequence accession numbers

The phage genomes assembled *de novo* were deposited in GenBank under accession numbers: MT670417.1 (ΦPsa267), MT670418.1 (ΦPsa300), MT670419.1 (ΦPsa315), MT670420.1 (ΦPsa347), MT670421.1 (ΦPsa381), MT670422.1 (ΦPsa397). The updated annotation of ΦPsa374 was deposited under accession number MT670055. The nucleotide sequences and the raw sequencing reads were uploaded to the NCBI SRA under the BioProject number PRJNA236447.

### Phylogenetic tree and phage genome comparison

Phages related to the ΦPsa374-like group were identified with Pairwise Sequence Comparison (PASC) (Bao *et al*., 2014). A phylogenetic tree was built with the Virus Classification and Tree building Online Resource (VICTOR) (Meier-Kolthoff and Göker, 2017) using distance formula *d*_6_ and branch support was inferred from 100 pseudo-bootstrap replicates each and edited with FigTree 1.4.3. Phage taxonomic analysis was performed in accordance to the International Committee on Taxonomy of Viruses (ICTV) and NCBI Taxonomy Database classifications. Whole genome comparison between ΦPsa374-like phages and distantly related phages was performed with blastn and tblastx in EasyFig 2.2.2 respectively. Multiple genome alignment of the tRNA region of the ΦPsa374-like phages was performed with MAFFT 7.450 (Katoh, 2002) using default settings. To identify close tRNA homologues, a BLASTN (Zhang *et al*., 2000) 2.11.0+ search (e-value<1E-5 cut-off) was used.

### Phage purification

To purify phage for EM and proteomic analysis 200 μL of overnight culture of *Psa* (OD_600_ ~2.5) was used to inoculate 20 mL of LB and incubated at 28°C shaking at 160 RPM until it reached exponential phase (OD_600_=0.3, ~5 × 10^8^ CFU/mL). The culture was infected with ΦPsa374 at a multiplicity of infection (MOI) of 0.01 and incubated for 5 h at 28°C shaking at 160 RPM to allow phage propagation. Next, NaCl was added to the cultures at a 4% w/v final concentration and incubated on ice for 1 h. Cell debris was removed by centrifugation at 11,000 *g* for 10 min at 4°C. The supernatant was decanted and PEG6000 was added (10% w/v final concentration), dissolved by stirring at room temperature and stored on ice overnight to precipitate the phage. The phage was pelleted by centrifugation at 11,000 *g* for 20 min at 4°C, the supernatant discarded, and the tubes dried by inversion for 5 min. The phage pellet was resuspended in 600 μL of phage buffer for 1 h at room temperature. The PEG and remaining cell debris were extracted from the phage suspension by adding chloroform (NaHCO_3_ saturated) in a 1:1 ratio, vortexing for 1 min and through a centrifugation step at 4,000 *g* for 15 min at 4°C. The aqueous phase containing phage was recovered, the phage was titrated, and protein concentration was determined using the BCA Protein Assay Kit (Thermo Fisher) following the manufacturer’s instructions. The phage sample was further purified with a CsCl gradient (1.33-1.7 g/mL). A gradient was prepared in a SW 32.1 tube and the phage sample was gently deposited on top. The tubes were centrifuged at 140,000 *g* for 3 h at 4°C. A defined white phage band was collected using a Pasteur pipette and dialysed (10,000 kDa) twice for 30 min and once overnight, against 250 volumes of distilled water to remove CsCl. Phages were concentrated by vacuum centrifugation for 2 h at medium temperature and the titre was determined and stored at 4°C.

### Proteomic analysis

To perform a proteomic analysis of the phage structural proteins, phage samples were prepared as described earlier (Frampton *et al*., 2015). Briefly, purified phage samples were mixed with 4 × SDS loading dye (40 mM Tris-HCl (pH 6.8), 40% glycerol, 4 mM EDTA, 2.5% SDS, 0.2 mg/L bromophenol blue), boiled for 5 min and separated on a 12% SDS-PAGE (polyacrylamide gel electrophoresis). The gel lane was fractionated into eight molecular weight fractions and subjected to in-gel digestion with trypsin and concentrated using a centrifugal vacuum concentrator. The peptides in each fraction were reconstituted in 15 μL of 5% (v/v) acetonitrile, 0.2% (v/v) formic acid in water and 5 μL were injected per run into an Ultimate 3000 nano-flow uHPLC-System (Dionex Co, Thermo Scientific) that was in-line coupled to the nanospray source of an LTQ-Orbitrap XL mass spectrometer (Thermo Scientific). Peptides were separated on an in-house packed emitter-tip column (75 um ID PicoTip fused silica tubing (New Objectives, Woburn, MA) filled with C-18 material (3 μm bead size) to a length of 12 cm).

The raw spectra obtained were processed through the Proteome Discoverer software (Thermo Scientific) using default settings to generate peak lists. Peak lists were then searched against a combined amino acid sequence database containing all ΦPsa374 sequence entries integrated into the full SwissProt/UniProt sequence database using the Sequest HT (Thermo Scientific), Mascot (www.matrixscience.com) and MS Amanda (http://ms.imp.ac.at/?goto=msamanda) search engines. The Percolator algorithm (http://per-colator.com) was used to estimate the False Discovery Rate (FDR). Peptide hits were filtered for a strict FDR of q < 0.01. Additional peptide score filters for each search engine were applied to eliminate very low scoring peptide hits that may have passed the Percolator FDR filter. Relative protein abundances were estimated by the averaged intensity of the three strongest unique peptide signals per protein using the TOP3 approach.(Silva *et al*., 2006)

### Transmission electron microscopy (TEM) and Cryo-Electron Microscopy (EM)

To examine phage by transmission electron microscopy (TEM), 10 μL of high titre phage stock (~10^9^ PFU/mL) was loaded onto plasma-glowed carbon coated 300 mesh copper grids (Electron Microscopy Sciences, Hatfield, PA, USA). After 60 s the excess specimen was removed by blotting and 10 μL of 1% w/v phosphotungstic acid (PTA) (pH 7.2) was applied to the grid to stain the samples and blotted off immediately. The grids were analysed in the Otago Micro and Nano Imaging (OMNI) facility and viewed in a Philips CM100 BioTWIN transmission electron microscope (Philips/FEI Corporation, Eindhoven, Holland) and images captured using a MegaView lll digital camera (Soft Imaging System GmbH, Münster, Germany).

To perform cryo-electron microscopy (EM), 3.5 μL of high titre phage samples were loaded onto plasma-glowed Quantifoil R2/2 holey carbon grids. The excess liquid was removed by blotting and the grid was flash frozen in liquid ethane using a Vitrobot Mark IV. Grids were viewed using a JEOL JEM-2200FS microscope operated at 200 kV and with an energy filter slit width of 22 eV. Image collection was controlled using the SerialEM software (Mastronarde, 2005). Zero-loss images at 25 fps at a total exposure of 1.5 s were captured onto a Direct Electron DE20 detector at an image pixel size of 2.3 Å and a total dose of 25 e-/Å^2^. A total of 365 38-frame movies each were aligned using Direct Electron’s frame alignment program.

For the capsid structure, images were imported into cisTEM (Grant *et al*., 2018) where CTF estimation was performed, particle picking and 2D classification. A total of 239 particles were selected to calculate an icosahedral reconstruction. The resolution was estimated to be 11.5 Å, using the gold standard Fourier Shell Ring correlation criterion with a 0.143 cutoff value. In order to investigate the structural features of the phage tail, only particles where the tail was clearly visible were selected. Image processing was done in Relion (Scheres, 2012). Regions corresponding to the neck regions and to the base plate were analysed separately. Based on 2D classification, the class averages presenting the clearest features were selected; for the neck region and base plate averages we summed 140 and 170 images, respectively. For the tail structure, we selected 750 tail regions and we used the helical reconstruction algorithms implemented in Relion (He and Scheres, 2017). The final reconstructions showed a helical rise of 37 Å and a twist of 25.5 degrees. The final map was filtered to a resolution of 20 Å. Figures were generated in Chimera (Goddard *et al*., 2018).

### Isolation of escape phage

Escape phages are mutants of the WT phage that have managed to overcome phage resistance. To isolate an escape phage that can infect *Psa* strain PI 345, a 100 μL aliquot of *Psa* strain PI 345 overnight culture (OD_600_ ~2.5) was used to seed 4 mL of molten soft LBA and poured onto an LBA plate. Serial ten-fold dilutions of a high titre ΦPsa374 phage stock (10^10^ PFU/mL) were spotted (10 μL) onto the soft overlay and the plate was incubated overnight at 28°C. Plaques in the spots with the highest phage concentration were picked and propagated overnight in a PI 345 culture (5 mL) at 28°C in shaking conditions (160 RPM). Purification of the escape phage was performed by plaque picking and passaging three times. Phage stocks were prepared as described earlier and stored at 4°C. PCR confirmation of SNPs in *gp080* and *gp081* was performed using Taq polymerase and the following primers: *gp080* (PF5915: GTGTCTGAACTCAATCCCTTTGTAGAAG and PF5916: TTATGTAATGAGGTTAGAGAGGAATCC) and *gp081* (PF4326: TGTGCTCTTCGAGAGGTCTCACCATGGATCTGATTAAATATGACATGGCGGATATCTG and PF4329: TGTGCTCTTCGCTTGGTCTCATACTTGGAAGTTCTGATCCACAGTGTGTG). PCR products were visualized on a 1.15 % w/v agarose gel supplemented with 500 ng/mL ethidium bromide. Gel bands of the expected size were purified using the Illustra GFX PCR DNA and Gel Band Purification kit according to the manufacturer’s instructions. Purified PCR products were Sanger-sequenced at Massey University, with 1 ng/100 bp of product mixed with 4 pmol of primer in a total volume of 20 μL.

### Phage and bacterial genome sequencing and SNP calling

To identify the mutations in the *Psa* plantlet isolates and the ΦPsa374 escape phage/mutant, whole genome sequencing was performed. Phage genomic DNA (gDNA) was extracted from a high titre phage stock (~10^10^ PFU/mL) and overnight bacterial cultures using the DNAeasy Blood and Tissue Kit (QIAGEN) following manufacturer’s instructions. DNA concentration was quantified by fluorimetry using the Qubit dsDNA HS Assay Kit and the Qubit Fluorometer (2.0) (Invitrogen) following the manufacturer’s instructions. Isolated gDNA was sent to the Massey Genome Service (Massey University, New Zealand) where libraries were prepared using the Nextera XT DNA Library Preparation Kit (Illumina), QC was checked with the Quant-iT dsDNA HS Assay for quantification and analysed using SolexaQA++, fastQC and fastQsceen. Sequencing was performed using Illumina MiSeq (2 x 150 bp) and the resulting reads were processed and trimmed using SolexaQA++ (v3.1.7.1). Phage genome assembly was performed using SPAdes 3.9 (Bankevich *et al*., 2012) and single nucleotide polymorphisms (SNPs) and mutations were detected by performing a whole genome alignment between ΦPsa374 and the escape phage with Geneious (v6.1.6) using default settings. For the plantlet bacterial isolates, reads were assembled *de novo* into contigs, and SNPs were identified by mapping reads against the *Psa* reference genome using Geneious (v6.1.6). SNPs were identified by running the ‘Find variation/SNPs’ option in Geneious (v6.1.6) with customized parameters at minimum coverage of 50 and a minimum of 90 % variant frequency. All SNPs identified in this work had >95 % variant frequency.

### Protein alignment

Protein alignment of the glycosyltransferase (locus tag IYO_025560 in *Psa*) in plantlet isolates was performed with Clustal Omega (Sievers *et al*., 2011) using default settings.

### Efficiency of plaquing (EOP) assays

To identify the host receptor for ΦPsa374 and the ΦPsa374-like phages, 10-fold dilutions of phage lysates were spotted in 10 μL droplets on soft overlay agar seeded with *Psa* transposon mutants (Mesarich *et al*., 2017) and plantlet isolates. Experiments were performed in biological triplicates, and phage titre was recorded in PFU/mL. Efficiency of plaquing (EOP) was calculated by dividing the phage titres determined on the transposon mutants or plantlet isolates by the average phage titre on WT *Psa* (ICMP 18884).

### Phage resistance infection time courses

Cultures of *Psa* and plantlet isolates were grown from an initial OD_600_ of 0.05 at 28°C, shaking (160 RPM) until reaching an OD_600_ of 0.2 (exponential phase). The cultures were diluted to an OD_600_ of 0.05, 180 μL aliquots were pipetted into a 96-well plate and 20 μL of diluted ΦPsa374 or escape phage lysates were added to produce an MOI=0.01. The 96-well plate was incubated in a Varioskan Flash plate reader (Thermo Scientific) at 28°C with 180 RPM shaking and OD_600_ measurements measured in 12 min increments for 15 h. Each condition was repeated in triplicate with data plotted as the mean plus or minus the SEM.

### Extraction and gel-based characterization of ΦPsa374-resistant *Psa* plantlet isolate and Tn5-mutant LPS

Cultures of *Psa* with Tn insertions in various LPS genes (Mesarich *et al*., 2017) and ΦPsa374-resistant isolates from this study were grown overnight in 2 mL LB supplemented with 50 μg/mL kanamycin (Km) where required. Cultures were centrifuged and resuspended in PBS to a standard OD_600_ of 2.0. 500 μL of each cell suspension was mixed with an equal volume of Tris-saturated phenol and extracted using the small-scale preparation described in Jayaraman *et al*. 2020 (Jayaraman *et al*., 2020). LPS were resolved using 4%–12% premade NuPAGE™ Bis-Tris SDS gels on a Novex system with MOPS as the running buffer as described by the manufacturer (Thermo-Fisher). Gels were stained with the Pro-Q™ Emerald 300 Lipopolysaccharide Gel Stain Kit (Molecular Probes, OR, USA).

### Pathogenicity plantlet assays

Plant infection for pathogenicity assays were carried out as described in Jayaraman *et al*. (Jayaraman *et al*., 2020). Briefly, *A. chinensis var. chinensis* ‘Hort16A’ plantlets, grown from axillary buds on Murashige and Skoog rooting medium without antibiotics in sterile 400 mL plastic tubs (“pottles”), were purchased (Multiflora, NZ). Plantlets were grown at 20°C under Gro-Lux fluorescent lights under long-day conditions (16 hr:8 hr, light:dark) and used when the plantlets were approximately 12 weeks old. Overnight LB liquid medium cultures of *Psa* were pelleted at 5,000 *g*, resuspended in 10 mM MgSO_4_, cell density determined by measuring the optical density at 600 nm, and reconstituted at OD_600_ = 0.05 (~10^6^ CFU/mL, determined by plating) in 500 mL of 10 mM MgSO_4_. Surfactant Silwet L-77 (Lehle Seeds) was added to the inoculum at 0.0025% (v/v) to facilitate leaf wetting. Pottles of ‘Hort16A’ plantlets were flooded with the inoculum, submerging the plantlets for 3 min, drained, sealed, and then incubated under previously described plant growth conditions.

*In planta* growth of *Psa* strains was assayed as described previously (McAtee *et al*., 2018). Briefly, leaf samples of four leaf discs per pseudobiological replicate, taken randomly with a 1-cm diameter cork-borer from three plants, were harvested at 2 h (day 0), day 6, and day 12 post-inoculation. All four replicates per treatment, per time point were taken from the same pottle. To estimate *Psa* growth inside the plant, the leaf discs were surface-sterilized, placed in Eppendorf tubes containing three sterile stainless-steel ball bearings and 350 μL 10 mM MgSO_4_, and macerated in a Storm 24 Bullet Blender (Next Advance) for two bursts of 1 min each at maximum speed. A 10-fold dilution series of the leaf homogenates was made in sterile 10 mM MgSO4 until a dilution of 10^-8^ and plated as 10 μL droplets on LB medium supplemented with nitrofurantoin and cephalexin. After 2 days of incubation at 20°C, the CFU per cm^2^ of leaf area was ascertained from dilutions.

### Phage preparation for plantlet assay

Phage lysates were prepared for plantlet assays using the following protocol. Firstly, 6 mL of a *Psa* 18884 overnight culture (OD_600_ ~2.5) was added to ~600 mL of LB in 2 L conical flasks and grown at 28°C with shaking (~180 RPM). Once these cultures reached an OD_600_ of ~0.2 (after 5 – 6 h), phage lysate was added to an MOI of 0.1, cultures were left static at room temperature for 20 – 30 min to allow phage adsorption. Cultures were then returned to incubators at 28°C with ~160 RPM shaking overnight (~18 h). Resultant cultures were then centrifuged at 6000 x g for 5 min to pellet bacterial debris. The supernatant containing phages was then transferred to Ti-45 ultracentrifuge tubes and centrifuged at ~210,000 *g* for 30 min to pellet the phages. The supernatant was decanted and the remaining phage pellets resuspended in phage buffer (10 mM Tris pH 7.4, 10 mM MgSO_4_, 0.01% w/v gelatine). Phage lysates were stored at 4°C until use.

### Plantlet assay

For each spray application of phage onto plantlets, 15 mL of high titre phage lysate (~10^10^-10^11^ PFU/mL) was diluted to 50 mL with phage buffer and then passed through a 0.22 μm syringe-driven filter. A sample of the phage lysate was diluted and plated to determine the titre (Figure S2). This 50 mL volume of lysate was sprayed onto plantlets in six pottles (Multiflora, NZ, more details in Pathogenicity plantlet assay methods section) until the plantlets were fully covered with phage lysate. Filtered phage buffer was used as a control for any effect of the spray application.

Fifty μL of a *Psa* 18884 overnight culture (OD_600_ ~2.5) was added to 50 mL of LB in a 250 mL conical flask and was grown at 28°C with ~160 RPM shaking overnight (~18 hours; OD_600_ ~2.5). The culture was centrifuged at 5752 *g* for 15 minutes to pellet the cells. The supernatant was removed and the cells resuspended in the same volume of 10 mM MgCl_2_ + 0.0025% v/v silwet. Five mL of washed cells were added to 495 mL of 10 mM MgCl_2_ + 0.0025% v/v silwet. This generated an inoculum with approximately 1×10^3^ CFU/mL. In a class II biosafety cabinet, the pottles of *A. chinensis var. chinesis* plantlets were flooded for 3 min with the *Psa* 18884 suspension. For each treatment condition (MgCl_2_ +silwet only, untreated *Psa*, phage only and phage treated *Psa*) pottles containing 5 plantlets were used, which represented a single biological replicate, biological triplicates of each condition were performed for the assay. The pottles were then drained and left to dry for 30 min. A mixture of 10 mM MgCl_2_ + 0.0025% (v/v) silwet was used to flood 3 negative control pottles to determine if flooding had any effect on the plantlets. Pottles that were flooded and sprayed with buffer were also included as negative controls to determine microbial background on plantlets. Plantlets were flooded with *Psa* on day 2 and sprayed with phage on days 1, 2, 4 and 8. Plantlet pottles were incubated in a growth cabinet (28°C, 70% humidity, 16:8 light dark cycle) for the duration of the experiment.

On each sampling day a leaf was removed from a plantlet in each pottle to determine bacterial and phage density. A100 mm^2^ disc of was taken from each plantlet leaf. Each leaf disc was put into a 2 mL screw cap tube containing 5 glass beads and 300 μL of 10 mM MgCl_2_. Leaf tissue was homogenized by bead beating for 1-2 min. *Psa* CFU counts were generated by taking 3 x 10 μL samples (technical triplicate) from the homogenate and making a 10-fold dilution series using 10 mM MgCl_2_ and plating 5 μL of each dilution on an LB agar plate. Phage PFU counts were generated by making a 10-fold dilution series of homogenised leaf material using phage buffer, 5 μL of each dilution was spotted onto an agar plate with a 0.35% soft-agar overlay containing 100 μL *Psa* overnight culture (OD_600_ ~2.5). *Psa* and phage ϕPsa374 samples were diluted and plated in technical triplicates for each biological replicate (leaves from separate pottles) on each sample day, with a detection limit of 60 CFU or PFU per leaf disc.

*Psa* plantlet isolates were picked from colonies used for the enumeration of bacterial density described above for the untreated and the phage treated conditions on each sample day. On earlier sample days (2, 3, 4 and 5) a smaller number of colonies or no colonies were noted for certain replicates therefore isolate numbers were fewer on these days. This resulted in a total number of 421 isolates, 263 were from untreated plantlets and 158 were from phage treated plantlets. These isolates were picked into 6 x 96-well 2 mL deep well plates with 1 mL of LB medium in each well and then cultured overnight, from these cultures glycerol stocks were made and stored at −80°C. Overnight cultures were grown from these *Psa* glycerol stock plates to an OD_600_ of ~2, diluted to an OD_600_ of ~0.05 and then screened for phage sensitivity or resistance. This utilized liquid infection assays described in the above methods section “Phage resistance infection time courses” using an MOI of ΦPsa374 between 0.005 – 0.01. ϕPsa374-resistant isolates from sample days 11 and 16 were further characterized as described in “Phage and bacterial genome sequencing and SNP calling” above.

## Acknowledgements

This work was supported by the Ministry of Business, Innovation and Employment Endeavour Fund and Zespri Group Ltd. L.M.M. was supported by a University of Otago Doctoral Scholarship, the University of Otago Postgraduate Publishing Bursary and the Division of Health Sciences Postdoctoral Fellowship. Thanks to Vivienne Young for ultracentrifuge advice, Sean Meaden for help with SNP and statistical analysis, Evelien Adraenssens for taxonomy advice and Erik Rikkerink (PFR) and Chandan Pal (Zespri) for comments on the manuscript.

