## Supplementary Information for "A lipopolysaccharide-dependent phage infects a pseudomonad phytopathogen and can evolve to evade phage resistance"

### Supplementary data

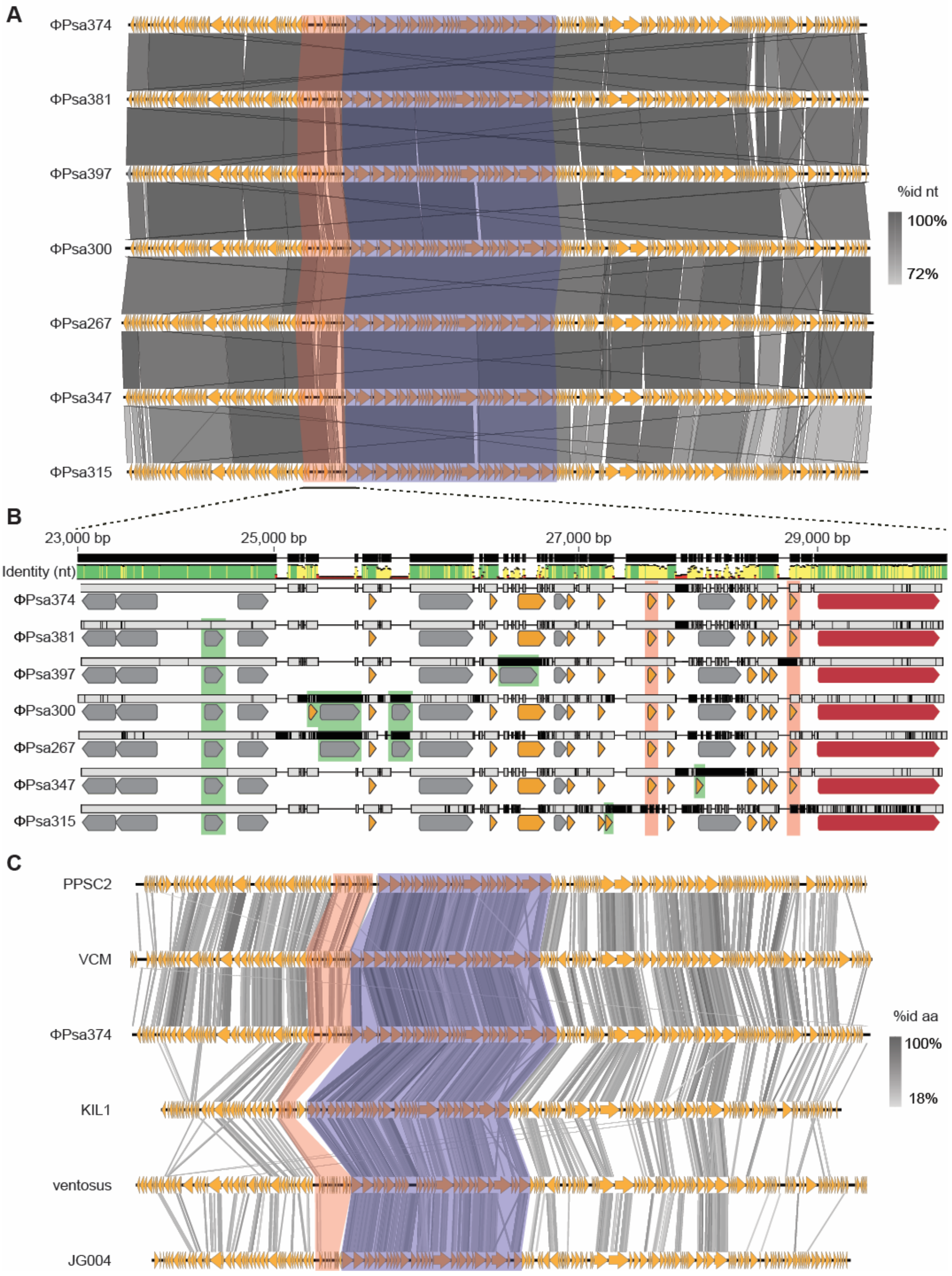


**Figure S1. The ΦPsa374-like phages have a high level of sequence conservation. A.** blastn whole genome alignment of the ΦPsa374 family tblastx whole genome alignment of phages distantly related to ΦPsa374. **B.** Sequence comparison of the region encoding tRNAs in ΦPsa374-like phages. Hypothetical proteins (grey), tail fiber (pink) and tRNAs (orange). Genetic features gained and lost are shown in green and red boxes, respectively. **C.** tblastx whole genome alignment of phages distantly related to ΦPsa374. In A. and C. conserved tRNAs and structural regions are highlighted in the orange and purple boxes, respectively.

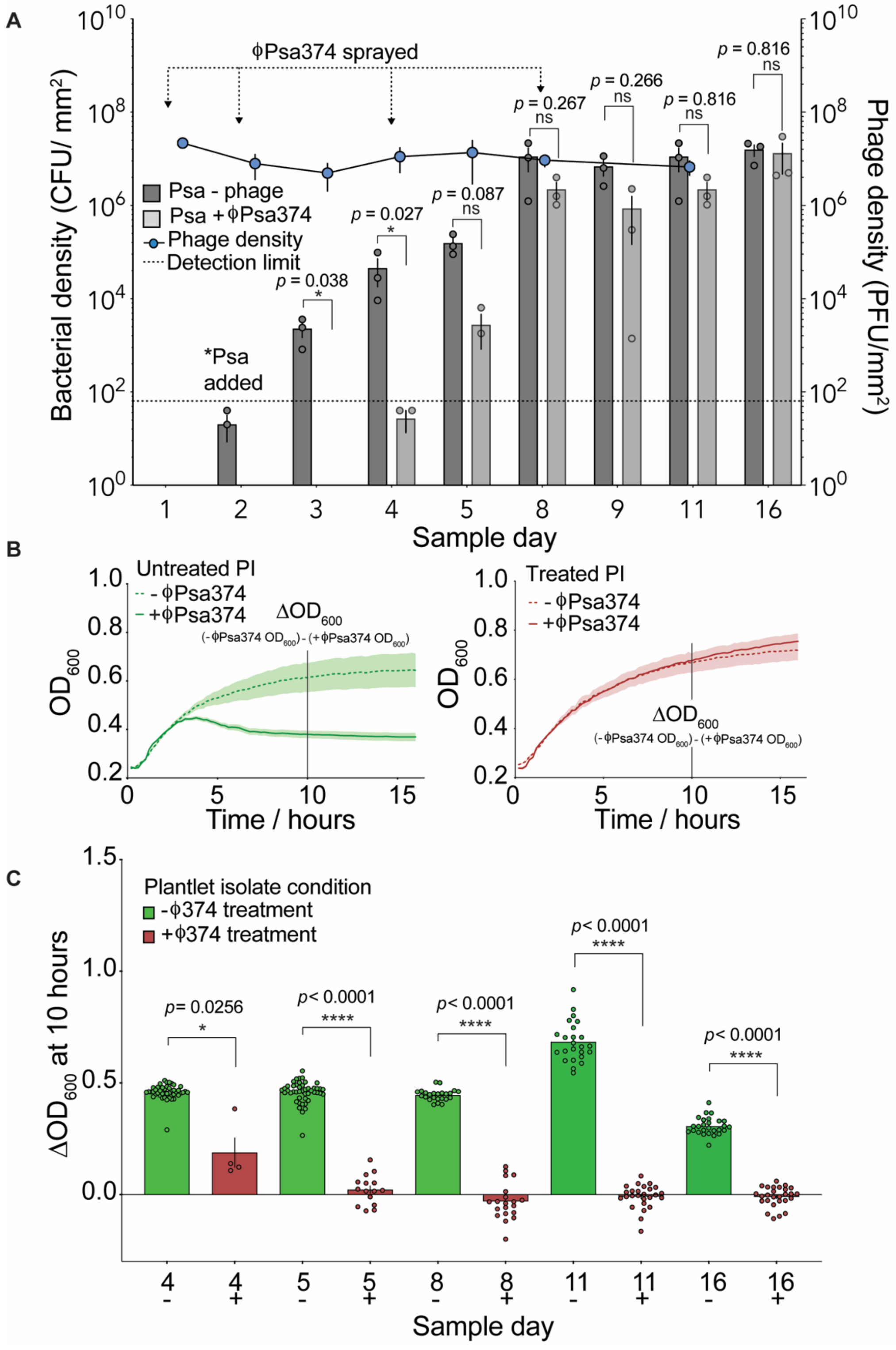


**Figure S2. Quadruple ΦPsa374 treatments of *Psa* infected kiwifruit plantlets reduces and delays *Psa* growth but results in phage resistance. A.** *Psa* and phage counts from a ϕPsa374 plantlet assay. *Actinidia chinensis* var. *chinensis* plantlets were sprayed with ϕPsa374 on day 1, 2, 4 and 8 and flooded with *Psa* on day 2. Leaf discs were removed from pottles at various time points used to enumerate bacteria (CFU/mm$^2$; grey bars) and phages (PFU/mm$^2$; blue circles). Each bar is the average and standard error of the mean (SEM) of three technical replicates repeated in biological triplicate. Each blue circle is the average and standard deviation of one plating replicate from three biological replicates. **B.** Examples of the growth ($OD_{600}$) response of *Psa* isolates from untreated and treated kiwifruit plantlets from sample day 16. Untreated plantlet isolates (PI) remain sensitive to ϕPsa374 indicated by a drop in $OD_{600}$ when ϕPsa374 is added to cultures. Treated PI continue to grow in a similar trend to the growth control when ϕPsa374 is added to cultures, indicating the development of resistance to ϕPsa374. **C.** Difference in $OD_{600}$ at 10 hours of growth between the *Psa* PI growth controls and inoculums with ϕPsa374 added from isolates from sample days 4, 5, 8, 11 and 16. A larger difference in $OD_{600}$ indicates that the *Psa* isolates remain sensitive to phage, while little to no difference indicates the development of resistance. *Psa* isolates from plantlets treated with ϕPsa374 were only able to be isolated from sample day 4 (n = 5), 80 % of these isolates show little to no difference in $OD_{600}$ indicating the development of ϕPsa374-resistance. From sample day 5 onwards all isolates show significant resistance to ϕPsa374. Technical and biological replicates are indicated by individual data points. For A. and C. unpaired Welch's t-tests were performed for each of the sample days between the mean of either the bacterial density or $\Delta OD_{600}$ between the untreated and ϕPsa374-treated conditions. Significance and *p* values are indicated for each data set.

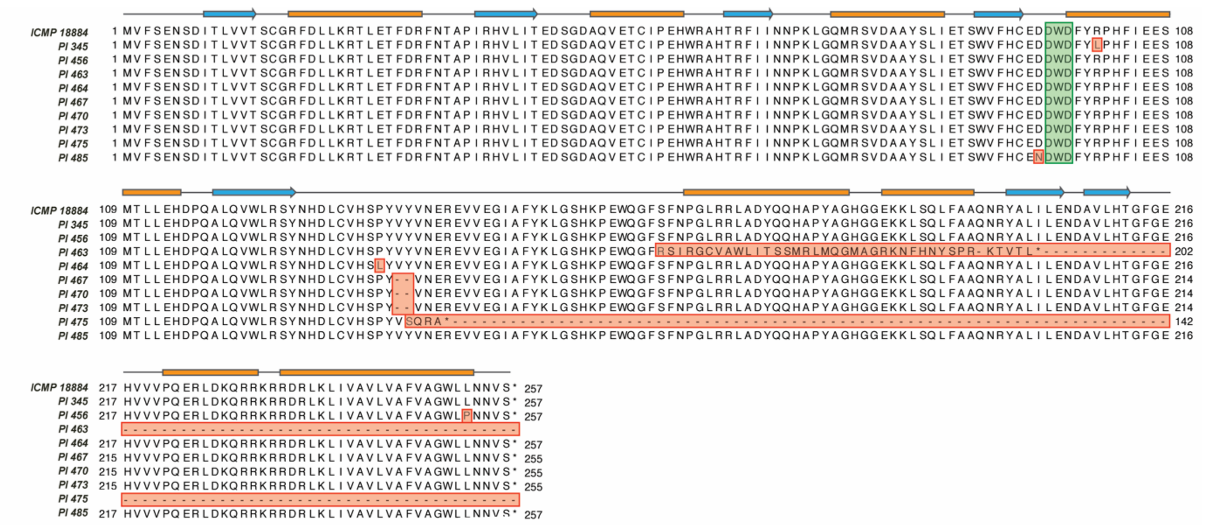


**Figure S3. ΦPsa374 treatment of *Psa* infected kiwifruit plantlets causes independent mutations in a glycosyltransferase.** Clustal Omega amino acid (aa) alignment of a glycosyltransferase found in the core LPS biosynthetic pathway (locus tag IYO_025560) of *Psa* isolates sampled from kiwifruit plantlets infected with *Psa* and treated with ΦPsa374. Mutations are highlighted in red and the DXD motif typical of this family of glycosyltransferases is highlighted in green. The Phyre2 predicted secondary structure is shown for the glycosyltransferase from *Psa* WT (ICMP 18884) with $\beta$-sheets represented by blue arrows and $\alpha$-helices by orange boxes.

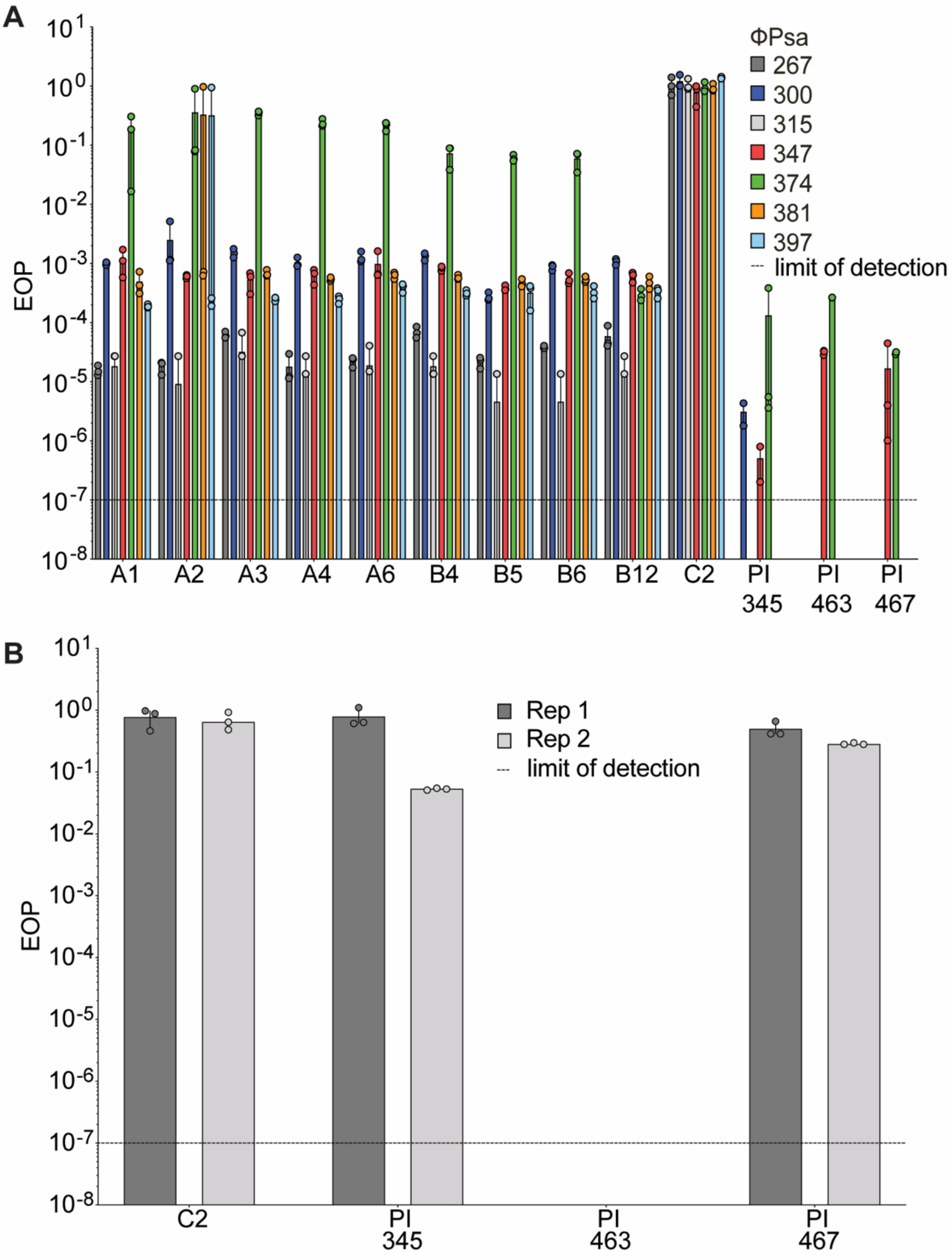


**Figure S4. ΦPsa374 and ΦPsa374-like phages utilize LPS as a receptor but can alter binding in response to LPS mutations. A.** Raw EOP data for ΦPsa374-like phage heat maps on a panel of LPS-Tn mutants (A1 – B12), a flagellum Tn mutant (C2) and three ΦPsa374-resistant plantlet isolates (Pl 345, 463 and 467) presented in Figure 3. **B.** EOP data for individual plaques picked and purified from a *Psa* Pl 345 lawn infected with ΦPsa374 against the flagellum Tn mutant (C2) and three ΦPsa374-resistant plantlet isolates (Pl 345, 463 and 467). Data for both A. and B. is presented in biological triplicate, each bar represents the mean while points represents each individual replicate. The limit of detection is $10^{-7}$, if no data point is present this indicates that plaques were not detected.

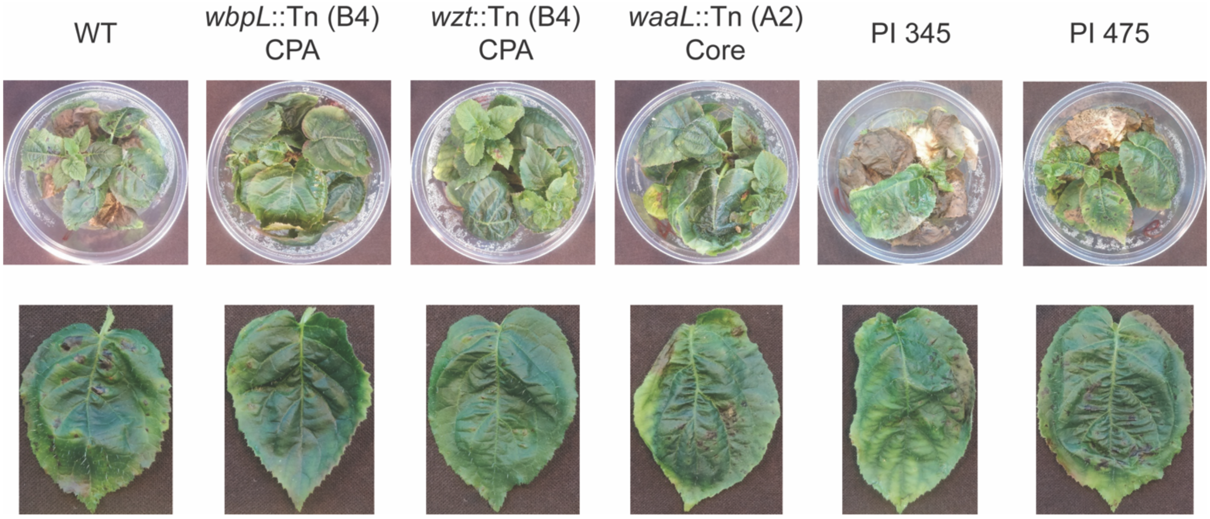


**Figure S5. ΦPsa374-resistant *Psa* from *in planta* pathogenicity studies retain pathogenicity, while LPS-Tn mutants do not.** Photographs of kiwifruit plantlets and representative leaves from *in planta* studies presented in Figure 4. A significant number of lesions associated with *Psa* disease can be seen in kiwifruit plantlets infected with WT, PI 345, PI 475. Kiwifruit plantlets infected with LPS-Tn mutants do not show any significant disease.

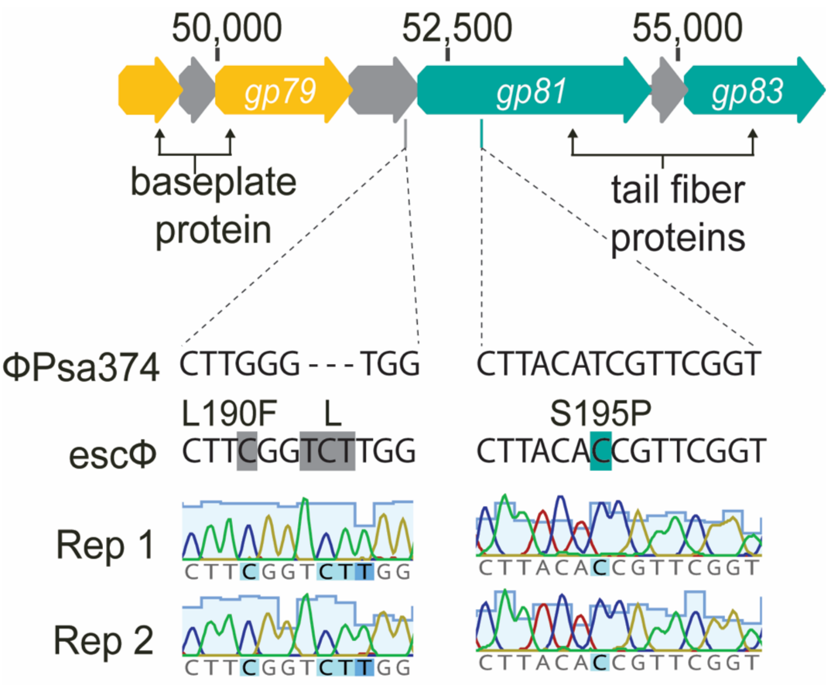


**Figure S6. PCR confirmation of mutations in *gp080* and *gp081* in ΦPsa374-escape phages able to overcome ΦPsa374-resistance in *Psa* plantlet isolates.** Two plaques (rep 1 and rep 2) found on a lawn of PI 345 infected with escΦ were picked, plaque purified and PCR amplification of genes *gp080* and *gp081* performed. Resultant bands were gel purified and sent for Sanger sequencing. Results for each plaque matched SNPs found in WGS.

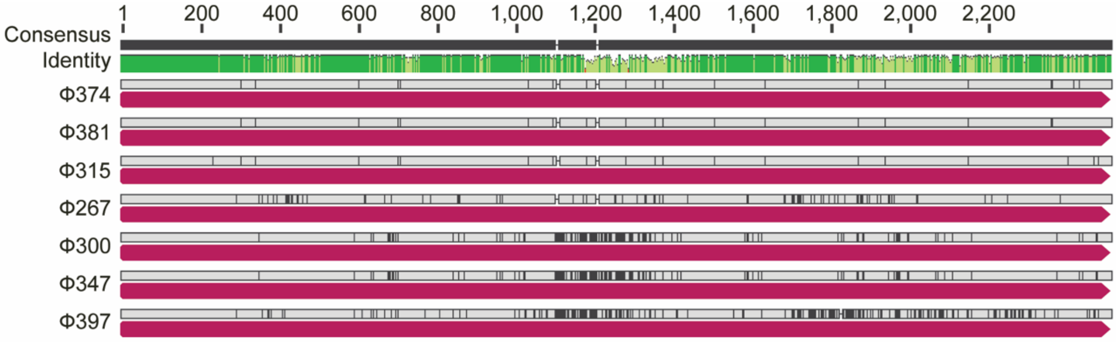


**Figure S7. Nucleotide alignment of *gp081* of ΦPsa374-like phages shows conservation at N-termini and variability at C-termini.** A multiple nucleotide alignment of *gp081* for ΦPsa374-like phages shows consensus identity conservation in dark green while areas of variability are indicated in light green. For each individual phage gene variation is highlighted in black above each CDS (magenta). Nucleotide alignment was performed in Geneious 6.1.6 using the Clustal Omega algorithm.

### Supplementary tables

#### **Table S1.** ΦPsa374 genome annotation

| ORFs | Name | Start (bp) | Stop (bp) | Direction |
|---|---|---|---|---|
| *gp001* | hypothetical protein CDS | 649 | 1122 | reverse |
| *gp002* | hypothetical protein CDS | 1318 | 1458 | reverse |
| *gp003* | hypothetical protein CDS | 1458 | 1655 | reverse |
| *gp004* | hypothetical protein CDS | 1655 | 2026 | reverse |
| *gp005* | hypothetical protein CDS | 2023 | 2145 | reverse |
| *gp006* | Peptide chain release factor 1 CDS | 2145 | 2537 | reverse |
| *gp007* | hypothetical protein CDS | 2530 | 2985 | reverse |
| *gp008* | hypothetical protein CDS | 3228 | 3470 | reverse |
| *gp009* | hypothetical protein CDS | 3636 | 4028 | reverse |
| *gp010* | hypothetical protein CDS | 4089 | 4433 | reverse |
| *gp011* | Tellurium resistance protein TerD CDS | 4493 | 5068 | reverse |
| *gp012* | Membrane protein TerC CDS | 5134 | 5805 | reverse |
| *gp013* | hypothetical protein with von Willebrand factor A-like domain TerF CDS | 5871 | 6512 | reverse |
| *gp014* | hypothetical protein CDS | 6528 | 6782 | reverse |
| *gp015* | Tellurite resistance protein TelA CDS | 6807 | 7907 | reverse |
| *gp016* | hypothetical protein CDS | 7912 | 8298 | reverse |
| *gp017* | hypothetical protein CDS | 8300 | 8803 | reverse |
| *gp018* | hypothetical protein CDS | 8800 | 9120 | reverse |
| *gp019* | hypothetical protein CDS | 9117 | 9320 | reverse |
| *gp020* | hypothetical protein CDS | 9329 | 9544 | reverse |
| *gp021* | DNA recombination-mediator protein A CDS | 9544 | 10122 | reverse |
| *gp022* | hypothetical protein CDS | 10122 | 10430 | reverse |
| *gp023* | hypothetical protein CDS | 10423 | 10584 | reverse |
| *gp024* | Putative phosphatase | 10565 | 11071 | reverse |
| *gp025* | hypothetical protein CDS | 11068 | 11190 | reverse |
| *gp026* | Nicotinamide phosphoribosyltransferase CDS | 11236 | 13011 | reverse |
| *gp027* | hypothetical protein CDS | 13077 | 13820 | reverse |
| *gp028* | hypothetical protein CDS | 13834 | 14238 | reverse |
| *gp029* | Ribose-phosphate pyrophosphokinase CDS | 14248 | 15135 | reverse |
| *gp030* | hypothetical protein CDS | 15198 | 15536 | reverse |
| *gp031* | hypothetical protein CDS | 15523 | 15729 | reverse |
| *gp032* | hypothetical protein CDS | 15729 | 16145 | reverse |
| *gp033* | RNA ligase, phage-associated CDS | 16145 | 17050 | reverse |
| *gp034* | hypothetical protein CDS | 17053 | 17274 | reverse |
| *gp035* | Phosphohydrolase (MutT/nudix family protein) CDS | 17274 | 17843 | reverse |
| *gp036* | Guanosine-3',5'-bis(diphosphate) 3'-pyrophosphohydrolase CDS | 17840 | 18310 | reverse |
| *gp037* | hypothetical protein CDS | 18368 | 18847 | reverse |
| *gp038* | DNA ligase, phage-associated CDS | 18858 | 20015 | reverse |
| *gp039* | hypothetical protein CDS | 20008 | 20247 | reverse |
| *gp040* | hypothetical protein CDS | 20244 | 20564 | reverse |
| *gp041* | hypothetical protein CDS | 20564 | 20797 | reverse |
| *gp042* | dCMP deaminase CDS | 20784 | 21188 | reverse |
| *gp043* | hypothetical protein CDS | 21379 | 21585 | reverse |
| *gp044* | hypothetical protein CDS | 21582 | 21926 | reverse |
| *gp045* | ATP-dependent Clp protease proteolytic subunit CDS | 21995 | 22606 | reverse |
| *gp046* | Phosphate starvation-inducible protein PhoH, predicted ATPase CDS | 22599 | 23360 | reverse |
| *gp047* | hypothetical protein CDS | 23402 | 23746 | reverse |
| *gp048* | hypothetical protein CDS | 23746 | 24186 | reverse |
| *gp049* | hypothetical protein CDS | 25033 | 25353 | forward |
| *tRNA* | tRNA_Val | 25821 | 25891 | forward |
| *gp050* | hypothetical protein CDS | 26137 | 26697 | forward |
| *tRNA* | tRNA_Trp | 26789 | 26862 | forward |
| *tRNA* | tRNA_Arg | 26959 | 27032 | forward |
| *gp051* | hypothetical protein CDS | 27118 | 27246 | forward |
| *tRNA* | tRNA_Glu | 27255 | 27328 | forward |
| *tRNA* | tRNA_Ile | 27553 | 27624 | forward |
| *tRNA* | tRNA_Asn | 27951 | 28033 | forward |
| *tRNA* | tRNA_Pro | 28155 | 28225 | forward |
| *gp052* | hypothetical protein CDS | 28462 | 28614 | forward |
| *tRNA* | tRNA_Leu | 28705 | 28778 | forward |
| *tRNA* | tRNA_Cys | 28839 | 28910 | forward |
| *tRNA* | tRNA_Gly | 28920 | 28992 | forward |
| *tRNA* | tRNA_Phe | 29001 | 29073 | forward |
| *gp053* | Phage tail fiber protein CDS – structural protein (MS result) | 29286 | 30563 | forward |
| *gp054* | hypothetical protein CDS | 30579 | 30842 | forward |
| *gp055* | Phage tail fiber protein CDS – structural protein (MS result) | 30842 | 32653 | forward |

| ORFs | Name | Start (bp) | Stop (bp) | Direction |
|---|---|---|---|---|
| *gp056* | hypothetical protein CDS | 32669 | 32896 | forward |
| *gp057* | Phage terminase, large subunit CDS | 33048 | 34517 | forward |
| *gp058* | Putative portal protein CDS – structural protein (MS result) | 34529 | 35986 | forward |
| *gp059* | Phage capsid and scaffold CDS | 35996 | 36451 | forward |
| *gp060* | hypothetical protein CDS | 36448 | 37368 | forward |
| *gp061* | Head decoration protein CDS – structural protein (MS result) | 37395 | 37781 | forward |
| *gp062* | Phage capsid protein CDS – structural protein (MS result) | 37794 | 38855 | forward |
| *gp063* | Phage protein CDS – structural protein (MS result) | 38906 | 39385 | forward |
| *gp064* | hypothetical protein CDS – structural protein (MS result) | 39369 | 39836 | forward |
| *gp065* | hypothetical protein CDS – structural protein (MS result) | 39848 | 40216 | forward |
| *gp066* | hypothetical protein CDS – structural protein (MS result) | 40213 | 40782 | forward |
| *gp067* | Structural protein CDS – structural protein (MS result) | 40794 | 42077 | forward |
| *gp068* | hypothetical protein CDS – structural protein (MS result) | 42112 | 42636 | forward |
| *gp069* | hypothetical protein CDS – structural protein (MS result) | 42701 | 43204 | forward |
| *gp070* | hypothetical protein CDS – structural protein (MS result) | 43214 | 43750 | forward |
| *gp071* | hypothetical protein CDS | 43770 | 44132 | forward |
| *gp072* | hypothetical protein CDS | 44153 | 44377 | forward |
| *gp073* | Tape measure protein CDS – structural protein (MS result) | 44390 | 46747 | forward |
| *gp074* | hypothetical protein CDS – structural protein (MS result) | 46747 | 47550 | forward |
| *gp075* | hypothetical protein CDS | 47561 | 47902 | forward |
| *gp076* | Putative structural protein CDS – structural protein (MS result) | 47906 | 48826 | forward |
| *gp077* | Putative baseplate protein CDS – structural protein (MS result) | 48823 | 49584 | forward |
| *gp078* | hypothetical protein CDS – structural protein (MS result) | 49593 | 49958 | forward |
| *gp079* | Putative baseplate protein CDS – structural protein (MS result) | 49961 | 51418 | forward |
| *gp080* | Phage protein CDS – structural protein (MS result) | 51430 | 52164 | forward |
| *gp081* | Phage tail fiber protein CDS – structural protein (MS result) | 52176 | 54653 | forward |
| *gp082* | hypothetical protein CDS | 54703 | 55071 | forward |
| *gp083* | Phage tail fiber protein CDS – structural protein (MS result) | 55090 | 56565 | forward |
| *gp084* | Putative phage-encoded peptidoglycan binding protein CDS | 56567 | 57124 | forward |
| *gp085* | hypothetical protein CDS | 57140 | 57343 | forward |
| *gp086* | hypothetical protein CDS | 57327 | 57749 | forward |
| *gp087* | hypothetical protein CDS | 57899 | 58198 | forward |
| *gp088* | hypothetical protein CDS | 58214 | 58540 | forward |
| *gp089* | hypothetical protein CDS | 58567 | 58755 | reverse |
| *gp090* | hypothetical protein CDS | 58748 | 58891 | reverse |
| *gp091* | hypothetical protein CDS | 58945 | 59298 | reverse |
| *gp092* | hypothetical protein CDS | 59364 | 59756 | reverse |
| *gp093* | Phage protein CDS | 60166 | 60474 | forward |
| *gp094* | putative DNA/pantothenate metabolism flavoprotein | 60471 | 61103 | forward |
| *gp095* | hypothetical protein CDS | 61100 | 61315 | forward |
| *gp096* | hypothetical protein CDS | 61312 | 61545 | forward |
| *gp097* | hypothetical protein CDS | 61547 | 61762 | forward |
| *gp098* | hypothetical protein CDS | 61759 | 61953 | forward |
| *gp099* | hypothetical protein CDS | 61950 | 62270 | forward |
| *gp100* | hypothetical protein CDS | 62331 | 62948 | forward |
| *gp101* | hypothetical protein CDS | 63352 | 63564 | forward |
| *gp102* | hypothetical protein CDS | 63548 | 63676 | forward |
| *gp103* | hypothetical protein CDS | 63673 | 63855 | forward |
| *gp104* | hypothetical protein CDS | 63855 | 64178 | forward |
| *gp105* | T7-like phage primase/helicase protein CDS | 64175 | 66025 | forward |
| *gp106* | Phage DNA polymerase I CDS | 66045 | 68369 | forward |
| *gp107* | hypothetical protein CDS | 68441 | 68719 | forward |
| *gp108* | hypothetical protein CDS | 68743 | 68913 | forward |
| *gp109* | hypothetical protein CDS | 68946 | 69359 | forward |
| *gp110* | hypothetical protein CDS | 69443 | 70462 | forward |
| *gp111* | hypothetical protein CDS | 70537 | 70803 | forward |
| *gp112* | hypothetical protein CDS | 70803 | 70916 | forward |
| *gp113* | hypothetical protein CDS | 70919 | 71497 | forward |
| *gp114* | hypothetical protein CDS | 71494 | 71769 | forward |
| *gp115* | Phage exonuclease CDS | 71810 | 72844 | forward |
| *gp116* | hypothetical protein CDS | 72844 | 73068 | forward |
| *gp117* | Putative HNH endonuclease CDS | 73065 | 73619 | forward |
| *gp118* | hypothetical protein CDS | 73616 | 73978 | forward |
| *gp119* | Putative phosphoesterase CDS | 73971 | 74753 | forward |
| *gp120* | hypothetical protein CDS | 74805 | 75023 | forward |
| *gp121* | hypothetical protein CDS | 75025 | 75243 | forward |
| *gp122* | hypothetical protein CDS | 75236 | 75436 | forward |
| *gp123* | Phage protein CDS | 75433 | 76188 | forward |
| *gp124* | Putative thymidylate synthase CDS | 76265 | 77206 | forward |
| *gp125* | hypothetical protein CDS | 77199 | 77537 | forward |
| *gp126* | Ribonucleotide reductase of class Ia, beta subunit CDS | 77546 | 78607 | forward |
| *gp127* | Ribonucleotide reductase of class Ia, alpha subunit CDS | 78600 | 80330 | forward |

| ORFs | Name | Start (bp) | Stop (bp) | Direction |
|---|---|---|---|---|
| *gp128* | hypothetical protein CDS | 80372 | 80728 | forward |
| *gp129* | hypothetical protein CDS | 80773 | 81054 | forward |
| *gp130* | hypothetical protein CDS | 81051 | 81290 | forward |
| *gp131* | hypothetical protein CDS | 81292 | 81426 | forward |
| *gp132* | hypothetical protein CDS | 81423 | 81623 | forward |
| *gp133* | hypothetical protein CDS | 81625 | 81990 | forward |
| *gp134* | hypothetical protein CDS | 81983 | 82138 | forward |
| *gp135* | hypothetical protein CDS | 82149 | 82376 | forward |
| *gp136* | hypothetical protein CDS | 82373 | 82570 | forward |
| *gp137* | hypothetical protein CDS | 82570 | 82779 | forward |
| *gp138* | Nucleoside Triphosphate Pyrophosphohydrolase CDS | 82790 | 83203 | forward |
| *gp139* | Chromosome partition protein smc CDS | 83200 | 83643 | forward |
| *gp140* | hypothetical protein CDS | 83709 | 83969 | forward |
| *gp141* | hypothetical protein CDS | 83988 | 84404 | forward |
| *gp142* | hypothetical protein CDS | 84404 | 84658 | forward |
| *gp143* | Phage protein CDS | 84739 | 85713 | forward |
| *gp144* | hypothetical protein CDS | 85713 | 85970 | forward |
| *gp145* | putative anti-sigma factor | 85970 | 86365 | forward |
| *gp146* | hypothetical protein CDS | 86362 | 86691 | forward |
| *gp147* | hypothetical protein CDS | 86695 | 86910 | forward |
| *gp148* | hypothetical protein CDS | 86910 | 87137 | forward |
| *gp149* | hypothetical protein CDS | 87143 | 87277 | forward |
| *gp150* | hypothetical protein CDS | 87352 | 87546 | forward |
| *gp151* | hypothetical protein CDS | 87663 | 88004 | forward |
| *gp152* | hypothetical protein CDS | 88001 | 89164 | forward |
| *gp153* | hypothetical protein CDS | 89157 | 89339 | forward |
| *gp154* | hypothetical protein CDS | 89539 | 89763 | forward |
| *gp155* | hypothetical protein CDS | 90290 | 91354 | forward |
| *gp156* | hypothetical protein CDS | 91428 | 91583 | forward |
| *gp157* | hypothetical protein CDS | 91586 | 91756 | forward |
| *gp158* | hypothetical protein CDS | 91853 | 92152 | forward |
| *gp159* | hypothetical protein CDS | 92304 | 92444 | forward |
| *gp160* | hypothetical protein CDS | 92623 | 93003 | forward |
| *gp161* | Phage protein CDS | 93068 | 93766 | forward |
| *gp162* | hypothetical protein CDS | 93822 | 94163 | forward |
| *gp163* | hypothetical protein CDS | 94272 | 94541 | forward |
| *gp164* | hypothetical protein CDS | 94568 | 95071 | forward |
| *gp165* | hypothetical protein CDS | 95144 | 95719 | forward |
| *gp166* | hypothetical protein CDS | 95719 | 95892 | forward |
| *gp167* | hypothetical protein CDS | 96061 | 96504 | forward |
| *gp168* | hypothetical protein CDS | 96569 | 96787 | forward |
| *gp169* | hypothetical protein CDS | 96856 | 97056 | forward |
| *gp170* | hypothetical protein CDS | 97056 | 97322 | forward |

**Table S2.** tRNAs encoded by Psa374-like phages.

| tRNAs | Psa347-like phage | | | | | | | Closest ortholog | | |
|---|---|---|---|---|---|---|---|---|---|---|
| | 374 | 381 | 397 | 300 | 267 | 347 | 315 | Organism | Identity (%) | e-value |
| **Ser** | - | - | - | + | - | - | - | *Pseudomonas* phage PPSC2 | 83.3 | 1.79E-14 |
| **Val** | + | + | + | + | + | + | + | *Pseudomonas* phage VCM | 88.7 | 1.02E-15 |
| **Trp** | + | + | + | + | + | + | + | *Pseudomonas* phage VCM | 8.7 | 1.53E-07 |
| **Arg** | + | + | - | + | + | + | + | *Pseudomonas* phage VCM | 91.9 | 5.01E-20 |
| **Glu** | + | + | + | + | + | + | + | *Pseudomonas* phage PPSC2 | 96.0 | 3.20E-13 |
| **Ile** | + | + | + | + | + | + | + | no match found | | |
| **Arg** | - | - | - | - | - | - | + | *Pseudomonas* phage VCM | 100 | 1.45E-26 |
| **Asn** | + | + | + | + | + | + | - | no match found | | |
| **Pro** | + | + | + | + | + | + | + | *Pseudomonas* phage VCM | 100 | 1.41E-26 |
| **Leu** | + | + | + | + | + | + | + | no match found | | |
| **Cys** | + | + | + | + | + | + | + | *Pseudomonas* phage PPSC2 | 91.4 | 4.5E-11 |
| **Gly** | + | + | + | + | + | + | + | *Pseudomonas* phage ventosus | 94.5 | 3.29E-22 |
| **Phe** | + | + | - | + | + | + | - | *Pseudomonas* phage PPSC2 | 78.1 | 6.35E-06 |
| **Total** | 11 | 11 | 9 | 12 | 11 | 11 | 10 | | | |

**Table S3.** Gene and ORF differences in the structural genome between ΦPsa374 and analogous *Pseudomonas* phages.

| Psaϕ374 | | | Orthologous phage structural proteins | | | | | | | | | |
|---|---|---|---|---|---|---|---|---|---|---|---|---|
| **Locus tag** | **Predicted function** | **Protein size / kDa** | **KIL3** [1] | **Protein size / kDa** | **KIL5** [1] | **Protein size / kDa** | **PaP1** [2] | **Protein size / kDa** | **PAK_P3** [3] | **Protein size / kDa** | **PaoP5** [4] | **Protein size / kDa** |
| *gp053* | Phage tail fiber protein | 44.7 | - | - | - | - | - | - | - | - | - | - |
| *gp054* | Hypothetical protein | 8.8 | *ORF41* (vB_PsyM_KIL3_0046) | 9.1 | - | - | - | - | - | - | - | - |
| *gp055* | Phage tail fiber protein | 63.0 | *ORF42* (vB_PsyM_KIL3_0047) | 30.58 | *ORF42* (vB_PsyM_KIL5_0051) | 30.62 | - | - | - | - | - | - |
| *gp056* | Hypothetical protein | 7.6 | *ORF43* (vB_PsyM_KIL3_0048) | 4.2 | *ORF43* (vB_PsyM_KIL5_0052) | 4.2 | - | - | - | - | - | - |
| *gp057* | Phage terminase, large subunit | 55.9 | *ORF44* (vB_PsyM_KIL3_0049) | 51.97 | *ORF44* (vB_PsyM_KIL5_0053) | 52.22 | *gp046* | 57.1 | *gp001* | *53.6* | *gp052* | 57 |
| - | - | - | *ORF45* (vB_PsyM_KIL3_0050) | 21.91 | *ORF45* (vB_PsyM_KIL5_0054) | 20.7 | - | - | - | - | - | - |
| *gp058* | Putative portal protein | 54.5 | *ORF46* (vB_PsyM_KIL3_0051) | 54.78 | *ORF46* (vB_PsyM_KIL5_0055) | 54.76 | *gp047* | 54.23 | *gp002* | 56.3 | *gp053* | 54.2 |
| *gp059* | Phage capsid and scaffold | 16.8 | *ORF47* (vB_PsyM_KIL3_0052) Putative methyltransferase | 17.1 | *ORF47* (vB_PsyM_KIL5_0056) Putative phage protease XkdF | 18.45 | *gp048* DNA-specific methyltransferase | 17.2 | *gp003* Putative methyltransferase | 15.2 | *gp054* | 17.1 |
| *gp060* | Hypothetical protein | 33.2 | *ORF48* (vB_PsyM_KIL3_0053) | 35.57 | *ORF48* (vB_PsyM_KIL5_0057) | 35.59 | *gp049* | 33.1 | *gp004* | 31.8 | *gp055* | 31 |
| *gp061* | Head decoration protein | 13.7 | *ORF49* (vB_PsyM_KIL3_0054) | 14.57 | *ORF49* (vB_PsyM_KIL5_0058) | 14.57 | *gp050* | 14.87 | *gp005* | 13.5 | *gp056* | 15 |
| *gp062* | Phage capsid protein | 39.5 | *ORF50* (vB_PsyM_KIL3_0055) | 39.84 | ORF50 (vB_PsyM_KIL5_0059) | 39.87 | *gp051* | 39.38 | *gp006* | 40.2 | *gp057* | 39 |

| Psaφ374 | | | Orthologous phage structural proteins | | | | | | | | | |
|---|---|---|---|---|---|---|---|---|---|---|---|---|
| **Locus tag** | **Predicted function** | **Protein size / kDa** | **KIL3** [1] | **Protein size / kDa** | **KIL5** [1] | **Protein size / kDa** | **PaP1** [2] | **Protein size / kDa** | **PAK_P3** [3] | **Protein size / kDa** | **PaoP5** [4] | **Protein size / kDa** |
| *gp063* | Phage protein | 18.0 | *ORF51* (vB_PsyM_KIL3_0056) | 23.06 | *ORF51* (vB_PsyM_KIL5_0060) | 22.6 | *gp052* | 18.1 | *gp007* | 18.6 | *gp059* | 18.1 |
| *gp064* | Hypothetical protein | 17.1 | *ORF52* (vB_PsyM_KIL3_0057) | 24.06 | *ORF52* (vB_PsyM_KIL5_0061) Putative RNA polymerase | 17.3 | *gp053* *Putative RNA polymerase* | 10.5 | *gp008* | 17.5 | *gp060* Putative RNA polymerase | 8.8 |
| *gp065* | Hypothetical protein | 13.9 | *ORF53* (vB_PsyM_KIL3_0058) | 13.97 | *ORF53* (vB_PsyM_KIL5_0062) | 13.97 | *gp054* | 14.3 | *gp009* | 14.3 | *gp061* | 14.3 |
| - | - | - | *ORF54* (vB_PsyM_KIL3_0059) | 28.64 | - | - | - | - | - | - | - | - |
| *gp066* | Hypothetical protein | 21.7 | *ORF55* (vB_PsyM_KIL3_0060) | 23.06 | *ORF54* (vB_PsyM_KIL5_0063) | 22.26 | *gp055* | 21.29 | *gp010* | 23.4 | *gp062* | 21.3 |
| - | Hypothetical protein | - | *ORF56* (vB_PsyM_KIL3_0061) | 25.9 | *ORF55* (vB_PsyM_KIL5_0064) | 8.6 | - | - | - | - | - | - |
| *gp067* | Structural protein | 45.8 | *ORF57* (vB_PsyM_KIL3_0062) | 45.76 | *ORF56* (vB_PsyM_KIL5_0065) | 45.76 | *gp056* | 46.37 | *gp011* | 47.9 | *gp063* | 47 |
| - | - | - | *ORF58* (vB_PsyM_KIL3_0063) Putative endonuclease | 26.7 | *ORF57* (vB_PsyM_KIL5_0066) Putative endonuclease | 11.4 | - | - | - | - | - | - |
| - | - | - | - | - | *ORF58* (vB_PsyM_KIL5_0067) Hypothetical protein | 4.0 | - | - | - | - | - | - |
| - | - | - | - | - | *ORF59* (vB_PsyM_KIL5_0068) Putative homing endonuclease | 7.3 | - | - | - | - | - | - |

| Psaϕ374 | | | Orthologous phage structural proteins | | | | | | | | | |
|---|---|---|---|---|---|---|---|---|---|---|---|---|
| **Locus tag** | **Predicted function** | **Protein size / kDa** | **KIL3** [1] | **Protein size / kDa** | **KIL5** [1] | **Protein size / kDa** | **PaP1** [2] | **Protein size / kDa** | **PAK_P3** [3] | **Protein size / kDa** | **PaoP5** [4] | **Protein size / kDa** |
| *gp068* | Hypothetical protein | 18.7 | *ORF59* (vB_PsyM_KIL3_0064) | 19.92 | *ORF60* (vB_PsyM_KIL5_0069) Putative structural protein | 18.81 | *gp057* Putative structural protein | 18.97 | *gp012* | 18.8 | *gp064* | 19 |
| *gp069* | Hypothetical protein | 18.0 | *ORF60* (vB_PsyM_KIL3_0065) | 18.0 | *ORF61* (vB_PsyM_KIL5_0070) | 22.93 | *gp058* | 18.4 | *gp013* | 18.3 | gp065 | 13.5 |
| *gp070* | Hypothetical protein | 19.0 | *ORF61* (vB_PsyM_KIL3_0066) | 19.14 | *ORF62* (vB_PsyM_KIL5_0071) | 19.15 | *gp059* Putative structural protein | 17.7 | *gp014* | 18.7 | gp066 | 17.7 |
| - | - | - | *ORF62* (vB_PsyM_KIL3_0067) Putative homing endonuclease | 12.9 | *ORF63* (vB_PsyM_KIL5_0072) Hypothetical protein | 7.8 | - | - | - | - | - | - |
| *gp071* | Hypothetical protein | 12.9 | *ORF63* (vB_PsyM_KIL3_0068) | 12.9 | *ORF64* (vB_PsyM_KIL5_0073) | 13.01 | *gp060* Putative structural protein | 13.6 | *gp015* | 13.9 | gp067 | 13.5 |
| *gp072* | Hypothetical protein | 8.7 | *ORF64* (vB_PsyM_KIL3_0069) | 11.1 | ORF65 (vB_PsyM_KIL5_0074) | 11.09 | *gp161* | 9.9 | *gp016* | 8.7 | - | - |
| *gp073* | Tape measure protein | 84.4 | *ORF65* (vB_PsyM_KIL3_0070) | 87.69 | *ORF66* (vB_PsyM_KIL5_0075) | 85.3 | *gp061* | 85.87 | *gp017* | 84.8 | *gp068* | 85.4 |
| *gp074* | Hypothetical protein | 29.4 | *ORF66* (vB_PsyM_KIL3_0071) | 29.54 | *ORF67* (vB_PsyM_KIL5_0076) | 29.86 | *gp062* | 28.5 | *gp018* | 27.9 | *gp069* | 28.5 |
| *gp075* | Hypothetical protein | 13.5 | *ORF67* (vB_PsyM_KIL3_0072) | 13.4 | *ORF68* (vB_PsyM_KIL5_0077) | 13.62 | *gp063* | 13.9 | *gp019* | 14.6 | *gp070* | 13.9 |
| *gp076* | Putative structural protein | 34.1 | *ORF68* (vB_PsyM_KIL3_0073) | 35.11 | *ORF69* (vB_PsyM_KIL5_0078) | 35.11 | *gp064* | 33.8 | *gp020* | 32.6 | *gp071* | 17.6 |

| Psaϕ374 | | | Orthologous phage structural proteins | | | | | | | | | |
|---|---|---|---|---|---|---|---|---|---|---|---|---|
| **Locus tag** | **Predicted function** | **Protein size / kDa** | **KIL3** [1] | **Protein size / kDa** | **KIL5** [1] | **Protein size / kDa** | **PaP1** [2] | **Protein size / kDa** | **PAK_P3** [3] | **Protein size / kDa** | **PaoP5** [4] | **Protein size / kDa** |
| *gp077* | Putative baseplate protein | 27.4 | *ORF69* (vB_PsyM_KIL3_0074) | 27.97 | *ORF70* (vB_PsyM_KIL5_0079) | 27.98 | *gp065* | 26.7 | *gp021* | 27.2 | *gp072* | 26.7 |
| *gp078* | Hypothetical protein | 13.4 | *ORF70* (vB_PsyM_KIL3_0075) Tail lysozyme | 13.3 | *ORF71* (vB_PsyM_KIL5_0080) Tail lysozyme | 16.41 | *gp066* | 14.2 | *gp022* | 14.1 | *gp073* | 14.2 |
| *gp079* | Putative baseplate protein | 51.7 | *ORF71* (vB_PsyM_KIL3_0076) | 53.46 | ORF72 (vB_PsyM_KIL5_0081) | 53.43 | *gp067* | 52.4 | *gp023* | 53.3 | *gp074* | 50 |
| *gp080* | Phage protein | 26.6 | *ORF72* (vB_PsyM_KIL3_0077) | 26.79 | *ORF73* (vB_PsyM_KIL5_0082) | 26.79 | *gp068* | 26.7 | *gp024* | 28.1 | *gp075* | 28 |
| *gp081* | Phage tail fiber protein | 85.5 | *ORF73* (vB_PsyM_KIL3_0078) | 72 | *ORF74* (vB_PsyM_KIL5_0083) | 72.03 | *gp069* | 69.79 | *gp025* | 70.2 | *gp076* | 74 |
| *gp082* | Hypothetical protein | 13.6 | *ORF74* (vB_PsyM_KIL3_0079) | 15.6 | *ORF75* (vB_PsyM_KIL5_0084) | 16.04 | *gp070* | 13.6 | *gp026* | 14.7 | *gp077* | 14.5 |
| *gp083* | Phage tail fiber protein | 51.7 | *ORF75* (vB_PsyM_KIL3_0080) | 51.18 | *ORF76* (vB_PsyM_KIL5_0085) | 51.17 | *gp071* | 53.09 | *gp027* | 52.9 | *gp078* | 55 |
| *gp084* | Endolysin | 20.6 | *ORF76* (vB_PsyM_KIL3_0081) | 20.84 | *ORF77* (vB_PsyM_KIL5_0086) | 20.84 | *gp072* | 20.9 | *gp028* | 20.8 | *gp079* | 20.8 |

*Cells shaded in orange indicate proteins identified by ESI-MS/MS, all protein sizes are predicted unless identified in ESI-MS/MS, dashes (-) indicate gene not present, annotations different to PsaΦ374 or orthologous phage are in red text.

**Table S4.** SNPs found in the gene IYO_025560 of *Psa* isolates sampled from ΦPsa374-treated kiwifruit plantlets infected with *Psa* ICMP 18884.

| Isolate | Sample day | Bio /tech. rep. no. | Mutation location | CDS position (bp) | Codon change | Protein effect |
|---|---|---|---|---|---|---|
| PI 345 | 11 | 3, 3 | 5,648,985 | 302 | CGT → CTT | Substitution |
| PI 456 | 16 | 1, 1 | 5,648,532 | 755 | CTC → CCC | Substitution |
| PI 463 | 16 | 1, 3 | 5,648,801 | 486 | AAC → A-C | Frame shift |
| PI 464 | 16 | 1, 3 | 5,648,883 | 404 | CCC → CTC | Substitution |
| PI 467 | 16 | 2, 1 | 5,648,877 | 405 | CCC, TAT, GTC → CCC | Deletion |
| PI 470 | 16 | 2, 1 | 5,648,877 | 405 | CCC, TAT, GTC → CCC | Deletion |
| PI 473 | 16 | 2, 2 | 5,648,877 | 405 | CCC, TAT, GTC → CCC | Deletion |
| PI 475 | 16 | 2, 2 | 5,648,875 | 409 | TAT → -A- | Frame shift |
| PI 485 | 16 | 3, 1 | 5,649,004 | 283 | GAT → AAT | Substitution |

**Table S5.** Strains used in this study.

| Strain | Notes | Reference |
|---|---|---|
| *Pseudomonas syringae* pv. *actinidiae* ICMP 18884 | Aka *Psa* (WT) | [5] |
| **Transposon mutants** | **Insertion** | **Reference** |
| A1 | gt1::Tn | [6] |
| A2 | *waaL*::Tn | [6] |
| A3 | *wbpL*::Tn | [6] |
| A4 | *rfbD*::Tn | [6] |
| A6 | methyltransferase::Tn | [6] |
| B4 | *wzt*::Tn | [6] |
| B5 | *wbpY*::Tn | [6] |
| B6 | methyltransferase::Tn | [6] |
| B12 | methyltransferase::Tn | [6] |
| C2 | *flgI*::Tn | [6] |
| **Plantlet isolates** | **Mutation*** | **Reference** |
| PI 406 (Control from plantlets not treated with phage at day 16) | N/A | This study |
| PI 345 | R101L | This study |
| PI 456 | L252P | This study |
| PI 463 | 1 bp deletion | This study |
| PI 464 | P135L | This study |
| PI 467 | 6 bp deletion | This study |
| PI 470 | 6 bp deletion | This study |
| PI 473 | 6 bp deletion | This study |
| PI 475 | 2 × 1 bp deletion | This study |
| PI 485 | D95N | This study |

*Locations of mutations in glycosyltransferase

**Table S6.** Phages used in this study.

| Phages | Reference |
|---|---|
| *Pseudomonas syringae* phage ΦPsa374 | [7] |
| *Pseudomonas syringae* phage ΦPsa267 | [7] |
| *Pseudomonas syringae* phage ΦPsa300 | [7] |
| *Pseudomonas syringae* phage ΦPsa315 | [7] |
| *Pseudomonas syringae* phage ΦPsa347 | [7] |
| *Pseudomonas syringae* phage ΦPsa381 | [7] |
| *Pseudomonas syringae* phage ΦPsa397 | [7] |
| *Pseudomonas syringae* phage ΦPsa21 | [7] |
| *Pseudomonas syringae* phage ΦPsa374 escape phage | This study |
